# Dehydration promotes intracellular lipid synthesis and accumulation

**DOI:** 10.1101/2025.08.07.669190

**Authors:** Joshua S. Carty, Jazlyn Selvasingh, Yvonne Zuchowski, Hyuck-Jin Nam, Clothilde Pénalva, Gayani Nanayakkara, Erin Q. Jennings, Kelsey Voss, Edith T. Adame, John T. Tossberg, Wei Sheng Yap, Meiling Melzer, Olga Viquez, A. Scott McCall, Elizabeth R Piotrowski, Ryoichi Bessho, Shirong Cao, Katrina L. Leaptrot, Alexandra C. Schrimpe-Rutledge, Simona G. Codreanu, Stacy D. Sherrod, John A. McLean, Jonathan B. Trapani, Matthew A. Cottam, Ma Wan, Deepti Shrivastava, Don A. Delker, Matthew H. Wilson, Clinton M. Hasenour, Louise Lantier, Irene Chernova, Jamey D. Young, Volker H. Haase, Jose Pablo Vazquez-Medina, Dylan K. Kosma, Peter Kim, Jean-Philippe Cartailler, Mingzhi Zhang, Roy Zent, Raymond C. Harris, Jason A. Watts, Andrew S. Terker, Fabian Bock, Jeffrey C. Rathmell, Aylin R. Rodan, Juan P. Arroyo

## Abstract

Lipids can be considered a water reservoir used to offset dehydration stress as their oxidation by the mitochondria generates water. However, whether dehydration and the ensuing hypertonic stress directly regulate lipid synthesis is unknown. We found that hypertonic stress decreases cellular oxygen consumption, increases intracellular lipid synthesis, and favors glutamine oxidation as a carbon precursor for lipid synthesis via remodeling mitochondrial metabolism. These findings provide a mechanism whereby cellular dehydration leads to intracellular lipid accumulation, functionally linking water availability to lipid storage.

## Introduction

Water is necessary for life and dehydration induced hypertonic stress is common to most life forms on earth. Consequently, organisms with lifelong limited access to water have developed mechanisms to generate water from lipids. However, it is currently unknown whether there is a direct mechanism through which hypertonic stress regulates lipid storage and breakdown.

Water that is generated within an organism is known as metabolic water. Marine mammals (e.g., *pinnipeds* and *cetaceans*) do not consume sea water, rather they rely on metabolic water production for the maintenance of water homeostasis ^1^. Similar observations on the reliance on metabolic water have been made in birds ^2^ and camels ^3^. Metabolic water is mainly generated as a byproduct of the oxidation of lipids, carbohydrates, and proteins in the mitochondria. Lipid oxidation is the most efficient precursor for metabolic water generation, as beta-oxidation yields about 100 g of water for every 100 g of palmitate that is oxidized. This is almost twice as much as the water produced by the oxidation of glucose or protein ^4^. Therefore, in the appropriate context, lipid accumulation represents water storage.

While much is known about lipid metabolism in the context of energy balance, the role of hypertonic stress in lipid metabolism is unclear ^5, 6, 7, 8, 9^. In plants, exposure to hypertonic stress can trigger the accumulation and subsequent breakdown of lipids ^6, 7, 10, 11, 12, 13, 14, 15^. Dehydration induced hypertonic stress results in an extracellular fluid tonicity that is higher than intracellular fluid tonicity. This in turn promotes intracellular water efflux, and compromises cell survival.

Therefore, the acute accumulation of lipids can provide cells a way to store water in a non-diffusible form that can later be accessed by the cell through lipid oxidation once extracellular and intracellular tonicity are matched. Thus, we hypothesized that hypertonic stress in the form of water restriction or exposure to a hypertonic medium triggers intracellular lipid accumulation. In this study, we demonstrate that hypertonic stress directly promotes intracellular lipid synthesis and remodels mitochondrial metabolism.

## Results

### Hypertonic stress promotes intracellular lipid synthesis and accumulation

The kidney, particularly the medulla, experiences wide variations in extracellular tonicity (−0.5-to-4-fold relative to serum osmolality, i.e. 150 mOsm/Kg – 1200 mOsm/Kg in humans) under physiologic conditions. Thus, the kidney is an ideal organ to understand the physiologic impact of hypertonic stress on cellular lipid metabolism. To test whether hypertonic stress promotes lipid accumulation, we treated kidney collecting duct (CD) cells with 100 mmol NaCl in lipid-free medium (to eliminate the possibility of increased uptake) and measured intracellular lipid droplet abundance. We found a significant increase in both the number and size of the lipid droplets in the NaCl treated collecting duct cells (Fig. 1A).

**Fig. 1.**
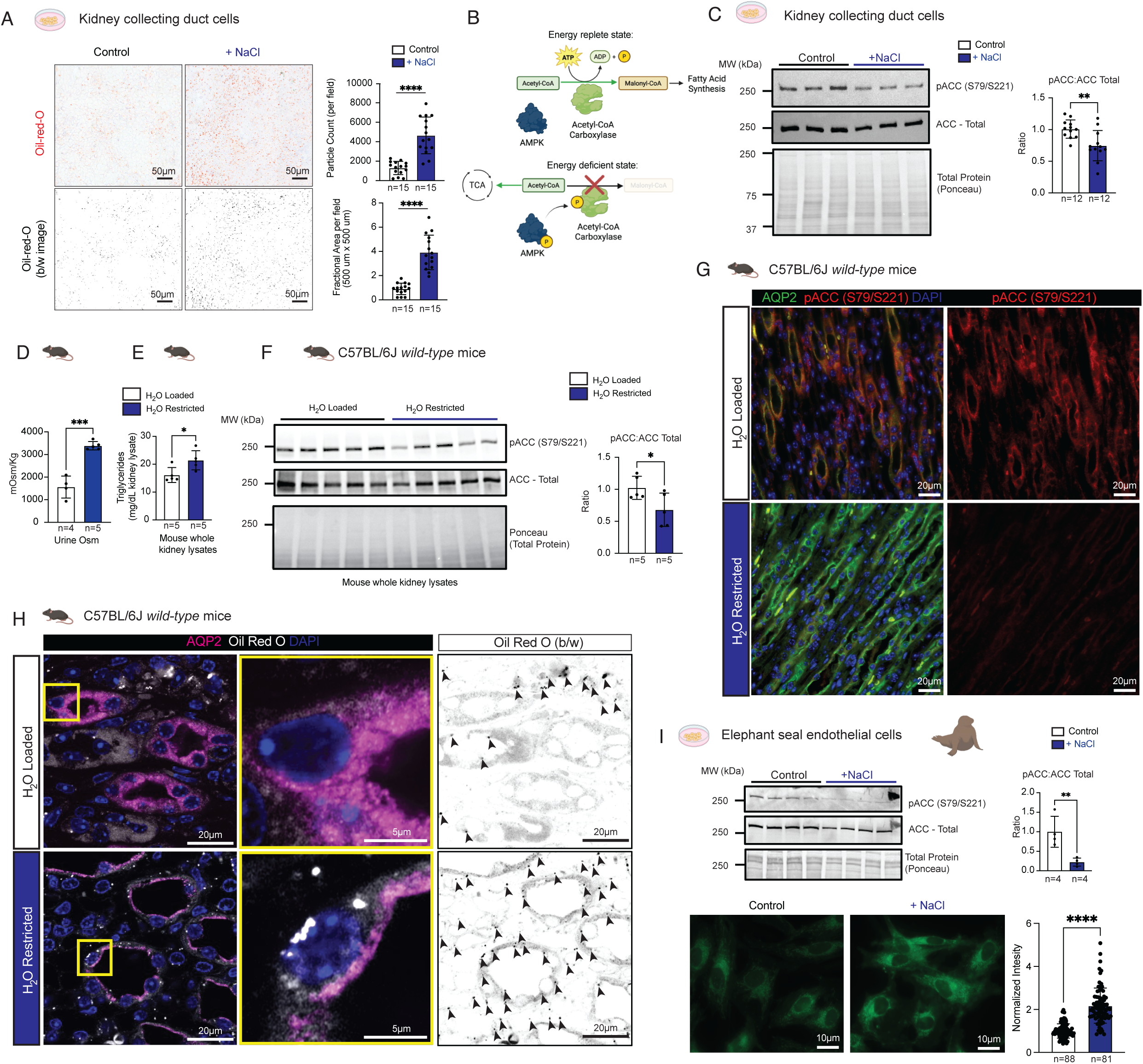
NaCl hypertonic stress induces lipid accumulation in cells. (A) Oil-red-o lipid droplet staining in CD cells treated with 100 mmol NaCl vs control 12 hours. (B) Acetyl-CoA Carboxylase is the rate limiting step in fatty acid synthesis and can be inhibited by AMP activated protein kinase (AMPK) via phosphorylation in states of energy depletion. (C) Immunoblots of total Acetyl-CoA Carboxylase (ACC) and pACC-S79/221 from kidney collecting duct cell lysates treated with 100 mmol NaCl vs control. (D) Urine osmolality (mOsm/Kg) and whole-kidney lysate triglyceride content from wild-type C57BL/6J mice after 24h water loading or water restriction. (F) Immunoblots of total ACC and pACC-S79/221 from whole kidney lysates of mice after water loading or water restriction. (G) Immunofluorescence of pACC-S79/221 and water channel Aquaporin 2 (AQP2), a collecting duct marker, in kidneys from water-loaded or water-restricted mice. (H) Oil-red-O lipid droplet and AQP2 immunofluorescent staining in kidneys from water-loaded and water-restricted mice. (I) Immunoblot of total Acetyl-CoA Carboxylase (ACC) and pACC-S79/221 and BODIPY lipid staining in from elephant seal primary endothelial cells lysates treated with 100 mmol NaCl vs control. Data presented as mean +/- SD. Data in A,C,D,E,F, and I analyzed with a two-tailed Student’s t-test. * p<0.05, ** p<0.01, *** p<0.001, ****p<0.0001, n as indicated in each panel. CD – Kidney collecting duct cells.

The rate limiting step for lipid synthesis is the activation of acetyl-coA carboxylase (ACC). Both ACC1 and ACC2 isoforms, which convert acetyl-coA to malonyl-coA that is used to synthesize fatty acids ^16, 17^, are expressed in the kidney ^18, 19^. They can be inhibited by AMP-activated protein kinase (AMPK) through phosphorylation of a carboxy-terminal serine residue, S79 and S221 in ACC1 and 2, respectively. Activating phosphorylation of AMPK during energy depletion favors phosphorylation (pACC-S79/221) and inhibition of ACC function, reducing lipid synthesis (Fig. 1B). Hypertonic NaCl stress in kidney CD cells resulted in ACC dephosphorylation and activation (Fig. 1C). 200 mM mannitol also dephosphorylated ACC, and incubation with isotonic control medium for 1 hr post 100 mmol NaCl reversed ACC phosphorylation, indicating the response is to hypertonic stress (fig. S1A,B). In parallel, the activating phosphorylation of AMPKα T172 (pAMPKα-T172) was decreased by hypertonic stress (fig. S1C). Thus, hypertonic stress inhibits AMPK, activates ACC, and favors intracellular lipid synthesis.

Next, we investigated if this response also occurred *in vivo*. Water restriction can increase murine kidney medullary interstitium tonicity to over 3500 mOsm/Kg (relative to the ∼300 mOsm/Kg in blood) to favor water reabsorption from the tubules into the blood. As water is retained, urine is concentrated and urine osmolality (UOsm) increases. We water restricted or water loaded wild-type C57BL/6J mice. We found that UOsm and whole kidney lysate triglyceride levels were higher in water-restricted mice (Fig. 1D, E). Water-restricted mice also had less pACC-S79/S221, consistent with increased ACC activation and lipid synthesis (Fig. 1F). In the collecting ducts of the water-restricted mice (identified by the water channel aquaporin 2, AQP2, which is upregulated and apically localized during water restriction ^20^), where the extracellular tonicity is highest, we saw near absent phosphorylation of pACC-S79/S221 (Fig. 1G) as well as increased intracellular lipid content (Fig. 1H). These results suggest hypertonic stress favors the intra-cellular accumulation of lipids both *in vitro* and *in vivo*.

To test whether this mechanism exists in animals that rely on metabolic water production, we used elephant seal (*Morounga angustirostris*) primary endothelial cells. When these cells were challenged with lipid-free NaCl stress we saw that pACC-S79/S221 decreased significantly and intracellular lipid increased, consistent with ACC activation and increased lipid synthesis (Fig. 1I). These results suggest that the effect of hypertonic stress on lipid accumulation can also occur in animals that rely on metabolic water.

Water restriction promotes fasting, which causes lipid accumulation in the kidney^21^. Thus, we carried out a pair-feeding experiment in which water loaded mice were fed a gelled diet which contained the same amount of chow consumed by the water restricted mice (Fig. 2A). Both groups of mice consumed the same number of calories (Fig. 2B), but the water restricted mice lost more weight (Fig. 2C), free fluid (Fig. 2D), andlean mass (Fig. 2E). but gained fat mass (2F). There were no significant differences in circulating triglycerides or free fatty acids (Fig. 2G, H). These results suggest water restriction can promote whole-body fat accumulation.

**Fig. 2.**
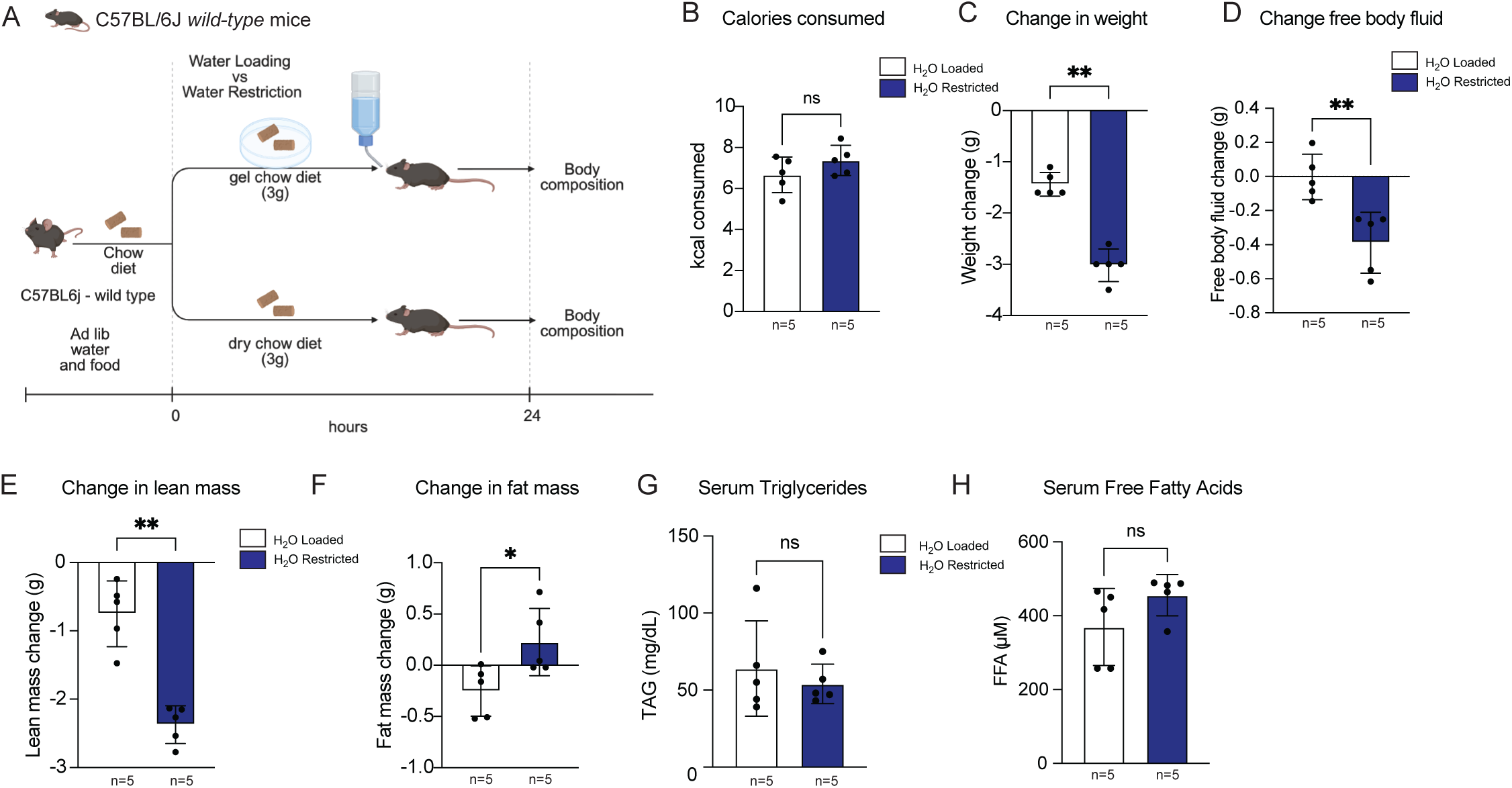
Water restriction increases fat mass in pair-fed wild-type mice. (A) Pair-fed C57BL6/J mice were water-restricted or water-loaded for 24 hrs (B) Calories consumed during-water restriction or water loading. Changes in weight (C), free body fluid mass (D), fat mass (E)and lean mass (F) after water restriction or water loading. Serum triglycerides (TAG) (G), and free fatty acids (H) after water restriction or water loading. Data is presented as mean +/- SD and analyzed with Mann-Whitney U test. * p<0.05, ** p<0.01, n as indicated in each panel. TAG (triglycerides), FFA (free fatty acids).

To further understand hypertonicity-induced lipid synthesis at the cellular level, we performed global untargeted lipidomic LC-MS/MS analysis to characterize the changes that occur after exposure to hypertonic stress in CD cells (fig. S2). We found a significant increase in overall length of all lipids analyzed (fig. S3A), and an increase in various lipid species with the most significant being ether-linked long-chain triacylglycerols and cardiolipins (Fig. 3A, fig. S2, S3), the latter of which are critical to maintaining mitochondrial membrane integrity ^22, 23^. In parallel, RNA-sequencing showed an enrichment in lipid synthetic pathways and a decrease in oxidative phosphorylation pathways in CD cells exposed to hypertonic stress (fig. S4A). Additionally, we found increased expression of *Fasn* and other targets of sterol regulatory element-binding protein 1c (SREBP-1c), a crucial transcription factor that regulates lipid accumulation (fig. S4B). Because may be more energetically favorable to prevent lipid breakdown rather than synthesis, we determined whether there were changes in the expression of enzymes responsible for lipid breakdown, adipose triglyceride lipase (ATGL) and hormone sensitive lipase (HSL). We found that mRNA for *Pnpla2* (which encodes for ATGL) was no different between groups (log2FC 0.0577, adjusted p-value=0.55263) while mRNA for *Lipe* (which encodes for HSL) increased log2FC 0.646 adjusted p-value =7.06×10^-14^, which argues against the inhibition of lipid breakdown. These data further suggest that hypertonic stress results in the upregulation of lipid synthesis *in vitro*.

**Fig. 3.**
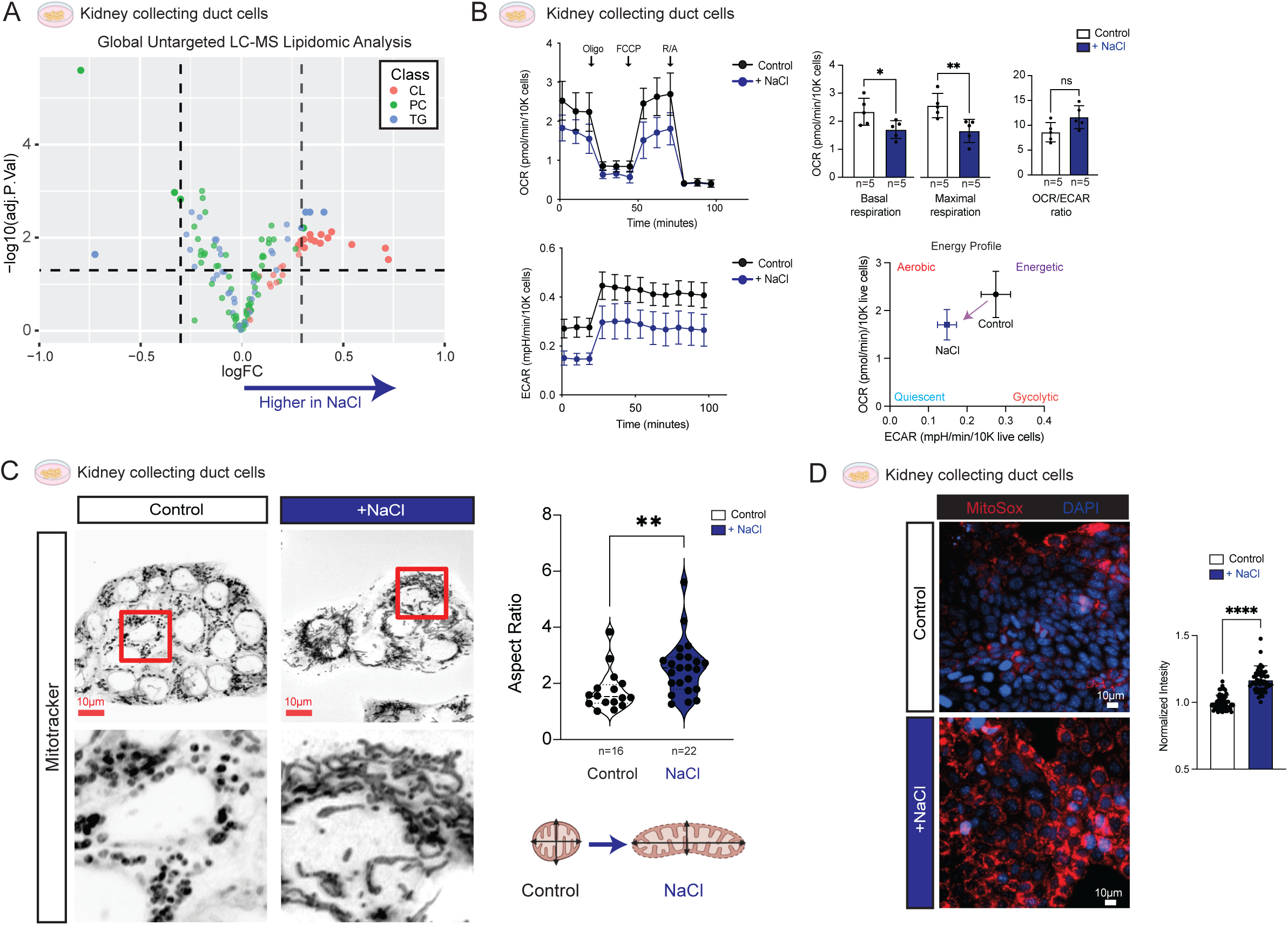
NaCl hypertonic stress alters mitochondrial metabolism. (A) Global untargeted LC-MS/MS lipidomic analysis of lipids by class in NaCl-treated CD cells vs controls. (B) Live metabolic cell profiling of NaCl treated CD cells. (C) Immunofluorescence mitochondrial aspect ratio analysis. Data in A LogFC of NaCl vs control cells plotted relative to -log10 (adj. p value). (D) Fluorescent staining of mitochondrial superoxide in NaCl-treated CD cells vs controls. Data in B and D presented as mean +/- SD and analyzed with Mann-Whitney test. Data in C presented as mean +/- SD box-violin plot and analyzed with a Mann-Whitney Test *p<0.05, ** p<0.01, n as indicated in each panel. CL - cardiolipin, TG - triglyceride, PC – Phosphatidylcholine, OCR – oxygen consumption rate, ECAR – extracellular acidification rate, CD – Kidney collecting duct cells.

To dissect the metabolic changes associated with hypertonic stress, we performed live cell metabolic profiling of CD cells. Salt-induced hypertonic stress resulted in decreased oxygen consumption rates (OCR) and decreased extracellular acidification (ECAR), indicating a shift toward a more quiescent energy profile (Figure 3B). To test if there were accompanying changes in mitochondrial morphology, we analyzed the mitochondrial aspect ratio of collecting duct cells treated with NaCl vs. controls. Hypertonic stress induced an increase in mitochondrial aspect ratio consistent with mitochondrial elongation, an adaptive strategy to maintain membrane potential and ATP production during cell stress (Fig. 3C) ^24, 25, 26^. These changes were accompanied by altered expression of mitochondrial fission/fusion related genes (fig. S4C). Additionally, hypertonic stress also caused mitochondrial elongation (fig. S5A) and decreased mitochondrial membrane potential (fig. S5B) in HeLa cells, in which decreased OCR with hypertonic stress was previously observed ^27^. Moreover, hypertonic stress induced increased mitochondrial superoxide (Fig. 3D) in line with decreased OCR and membrane potential. Together, these data suggest that hypertonic stress decreases mitochondrial metabolism and increases lipid synthesis simultaneously.

### Transcription factor NFAT5 is required for hypertonicity induced lipid accumulation

Nuclear factor of activated T-Cells 5 (NFAT5), also known as tonicity-responsive enhancer-binding protein (TonEBP), is a transcription factor that regulates the cellular response to hypertonic stress ^8^ and has been implicated in the regulation of fat mass ^28^. Total body knockout of *Nfat5* is embryonic lethal ^29^, but a haploinsufficient mouse model lacking one copy of *Nfat5* is protected from obesity when fed a high-fat diet, suggesting NFAT5 plays a critical role in lipid metabolism ^28^. We hypothesized that hypertonicity induced lipid synthesis requires NFAT5 activation. To test this we knocked down *Nfat5* in kidney collecting duct cells.We found that hypertonicity increased ACC activity by decreasing S79/221 phosphorylation, while *Nfat5* knockdown decreased ACC activity, increased AMPK activity, and decreased intracellular lipid synthesis and accumulation in NaCl-treated CD cells (Fig. 4 A,B fig. S6). To further ascertain if NFAT5 was involved in lipid synthesis, we performed Cleavage Under targets & Tagmentation (CUT&TAG) and measured NFAT5 occupancy of the regulatory regions of genes associated to lipid synthesis and elongation before and after hypertonic stress. The promoters for *Acly*, *Acaca*, *Fasn*, *Scd1*, *Elovl6*, and *Tecr* are marked by the active histone mark H3K27 acetyl under both isotonic and hypertonic conditions and show markedly increased NFAT5 occupancy following hypertonic stress (Fig. 4C). This suggests hypertonic stress induced activation of NFAT5 directly modifies the expression of genes related to lipid synthesis. To test if NFAT5 activation was required *in vivo* to promote lipid accumulation in the kidney, we used a haploinsufficient *Nfat5* knockout model, *Nfat5*^tm1a(KOMP)Wtsi^, herein referred to as *Nfat5* heterozygous KO (Fig. 4D) ^30^.

**Fig. 4.**
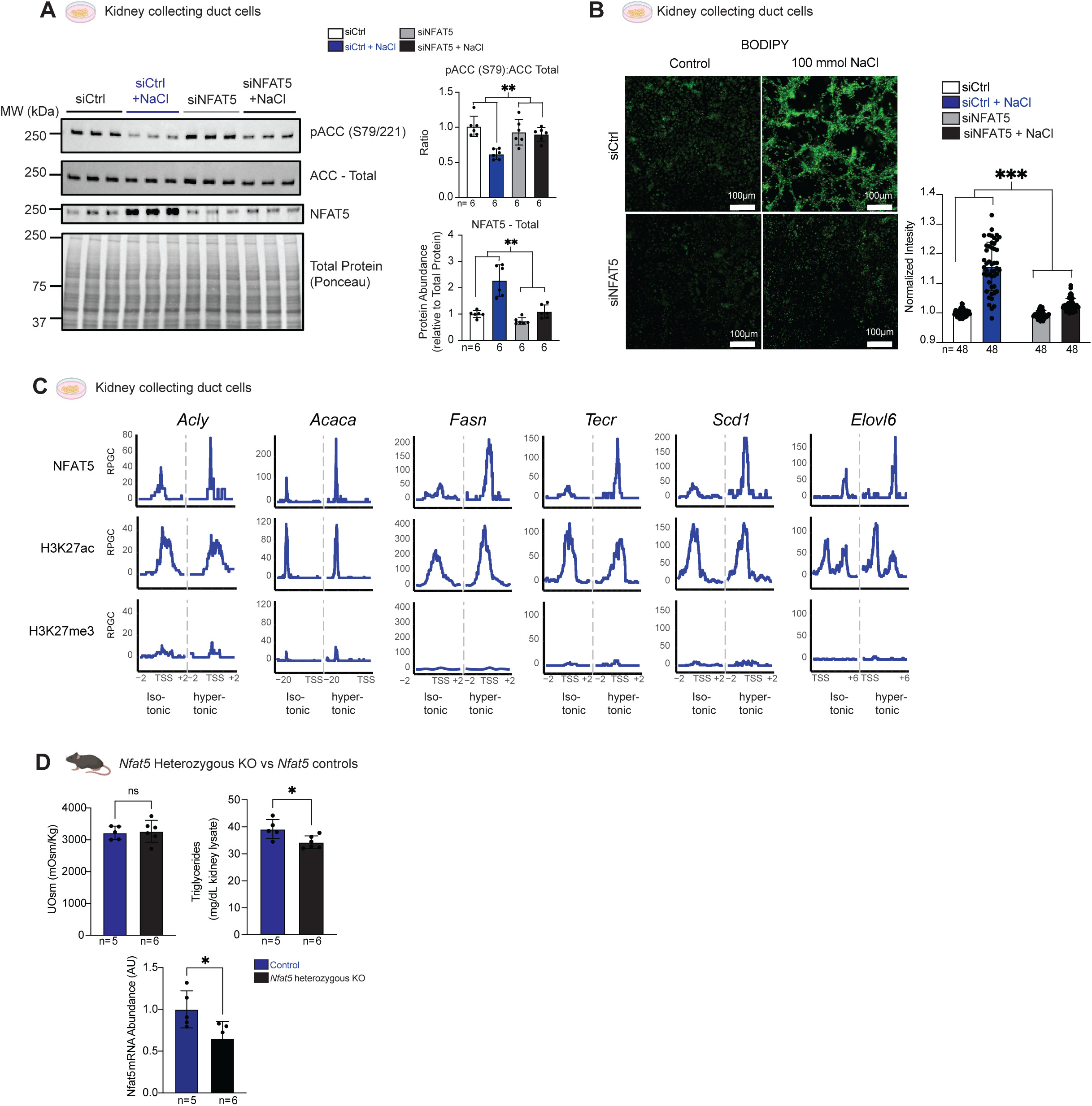
NFAT5 is required for hypertonicity induced lipid synthesis. (A) Immunoblots of total Acetyl-CoA Carboxylase (ACC), pACC-S79/221, and NFAT5 in CD cells transfected with siNfat5 vs siCtrl treated with NaCl or control. (B) Immunofluorescence of BODIPY-stained lipid droplets in CD cells treated with siCtrl or siNfat5 +/- NaCl. (C) Cleavage Under targets & Tagmentation (CUT&TAG) of kidney collecting duct (CD) cells under isotonic and hypertonic conditions showing NFAT5 occupancy in the promoter region of genes associated to lipid synthesis and/or elongation, H3K27ac (positive control), and H3K27me3 (negative control). y-axis denotes reads per genomic content (RPGC). (D) Urine osmolality, kidney lysate triglycerides, and Nfat5 mRNA in NFAT5 heterozygous KO mice vs controls after water restriction. Data in A and B presented as mean +/- SD, analyzed with two-way ANOVA with NaCl and siRNA as the independent variables. **p<0.01 for the interaction, n as noted in figure. Data in D presented as mean +/- SD analyzed with a two-tailed Student’s t-test *p<0.05. n as indicated in each panel. CD – Kidney collecting duct cells.

Following water restriction NFAT5 heterozygous KO mice had lower kidney lysate triglyceride content compared to wild-type controls, despite similar UOsm ^31^ (Fig. 4D). Together, these data show that NFAT5 is a critical component of hypertonicity-induced lipid synthesis and accumulation in cells and *in vivo*.

### Hypertonic stress-induced lipid synthesis requires glutamine oxidation

Lipid synthesis requires a carbon source. Glucose-derived pyruvate can be converted to acetyl-CoA via pyruvate dehydrogenase (PDH) and further metabolized by the tricarboxylic acid (TCA) cycle to citrate, a precursor for fatty acid synthesis (Fig. 5A). However, NaCl treatment of CD cells increased inhibitory phosphorylation of PDH serine 293, a site phosphorylated by pyruvate dehydrogenase kinase 1 (Fig. 5B), as also observed previously in HeLa cells exposed to hypertonic stress ^27^. Additionally, the medium glucose concentration was significantly higher in CD cells after treatment with NaCl vs. controls (Fig. 5C), suggesting decreased glucose consumption, which is in line with PDH inhibition and decreased ECAR (Fig. 3B) during hypertonic stress. These data suggest that there is an alternative carbon source for lipid synthesis.

**Fig. 5.**
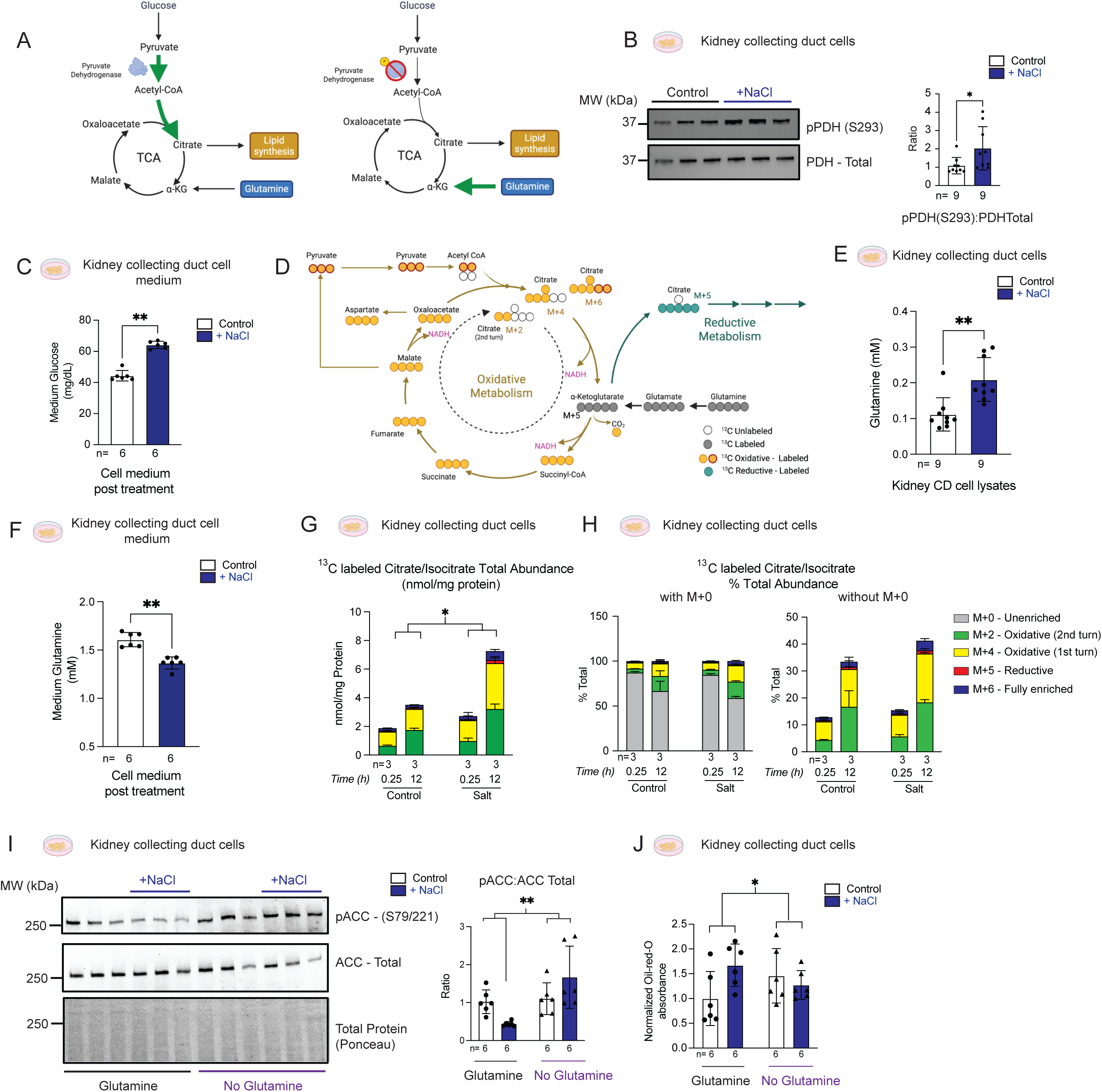
Glutamine is required for hypertonicity-induced lipid synthesis in vitro. (A) Pyruvate dehydrogenase (PDH) is inhibited via phosphorylation. (B) Immunoblotting for phosphorylated pPDH (S293) and total PDH in CD cells treated with NaCl vs control. (C) Cell medium glucose levels after treatment with NaCl vs control (D) Metabolic pathways of universally labeled 13C glutamine. (E) Intracellular glutamine levels in CD cells treated with NaCl vs control. (F) Glutamine levels in cell medium after treatment with NaCl vs control. (G) Total abundance of 13C labeled citrate/isocitrate isotopologues (nmol/mg protein) and (H) % distribution of 13C-enriched isotopologues of citrate/isocitrate in CD cells treated with control or NaCl hypertonic stress. (I) Immunoblotting for phosphorylated acetyl-coA carboxylase (pACC-S79/221), total ACC and normalized oil-red-o absorbance in kidney collecting duct cell lysates treated with NaCl vs control in presence or absence of glutamine. Data in B,C, E, and F presented as mean +/- SD, analyzed with a two-tailed student’s t-test *p<0.05, ** p<0.01. Data in G and H presented as mean +/- SD, analyzed with two-way ANOVA with NaCl and time as the independent variables *p<0.05 for the interaction in G, NS interaction in H. Data in I presented as mean +/- SD, analyzed with two-way ANOVA with NaCl and glutamine supplementation as the independent variables. *p<0.05, **p<0.01 for the interaction, n as noted in figure. CD – Kidney collecting duct cells.

The amino acid glutamine can be converted to the lipid precursor, citrate, via oxidative or reductive TCA cycling (Fig 5D) ^32, 33^ In cancer cells with decreased mitochondrial function, glutamine can be used as a precursor for lipid synthesis via reductive carboxylation ^32^. To test if hypertonic stress was mimicking a cancer cell-like phenotype, we first measured intracellular glutamine levels in CD cells exposed to hypertonic stress and found that intracellular glutamine increased (Fig 5E) and medium glutamine decreased (Fig. 5F) after exposure to NaCl, suggesting increased glutamine uptake by CD cells exposed to hypertonic stress. In fact, we found that mRNA expression for glutamine transporter SLC38A2 is increased in NaCl treated CD cells (log2FC 0.5376 adjusted p-value of 1.06*10^-50^). To understand whether the proportion of glutamine oxidation or reduction changed under hypertonic stress, we treated NaCl-exposed CD cells with uniformly labeled ^13^C glutamine. The abundance of carbon-13 (i.e., ^13^C-enriched) isotopologues of citrate/isocitrate M+2, M+4, M+5, and M+6 were elevated after salt treatment (Fig 5G). However, to our surprise, reductive carboxylation of α-ketoglutarate to citrate (M+5) was minimal in both NaCl-and control-treated collecting duct cells and no different between groups (Fig. 5H). Together, these results suggest that hypertonic stress does not promote the reductive carboxylation of glutamine, but promotes the incorporation of glutamine into the TCA cycle in the oxidative direction.

To further evaluate the role of glutamine in the hypertonic stress-induced lipid synthesis response ^32, 33^ we removed glutamine from the cell medium and exposed the CD cells to hypertonic stress. In cells exposed to hypertonic stress, glutamine removal resulted in inhibitory ACC phosphorylation (Fig. 5I) and decreased intracellular lipid content (Fig.5J, fig S7). Together, these results suggest that glutamine is critical to the lipogenic response to hypertonicity *in vitro*.

We then explored whether the requirement for glutamine in response to hypertonic stress is evolutionarily conserved and required for the *in vivo* response to hypertonic stress using a *Drosophila melanogaster* model. Flies were fed normal or high salt (0.3M added NaCl) for 24 hours, followed by 24 hours of water restriction (empty vial) or wet starvation (water without food) (Fig. S8). Glutamine carbons primarily enter the TCA cycle via glutaminase (GLS) conversion of glutamine to glutamate, and glutamate dehydrogenase (GDH) conversion of glutamate to α-ketoglutarate (Fig. 6A). *D. melanogaster* mutants in which *Gls* or *Gdh* were knocked down in the renal epithelium (Malpighian tubule) had significantly higher mortality in response to NaCl hypertonic stress (Fig. 6A, fig. S9A), as did wild-type flies fed the GLS inhibitor, BPTES (fig. S10B). Wet starvation had no effect on mortality (fig. S10C, S9C). Together these results show that glutamine incorporation into the TCA cycle is required to prevent hypertonic stress induced mortality *in vivo*.

**Fig. 6.**
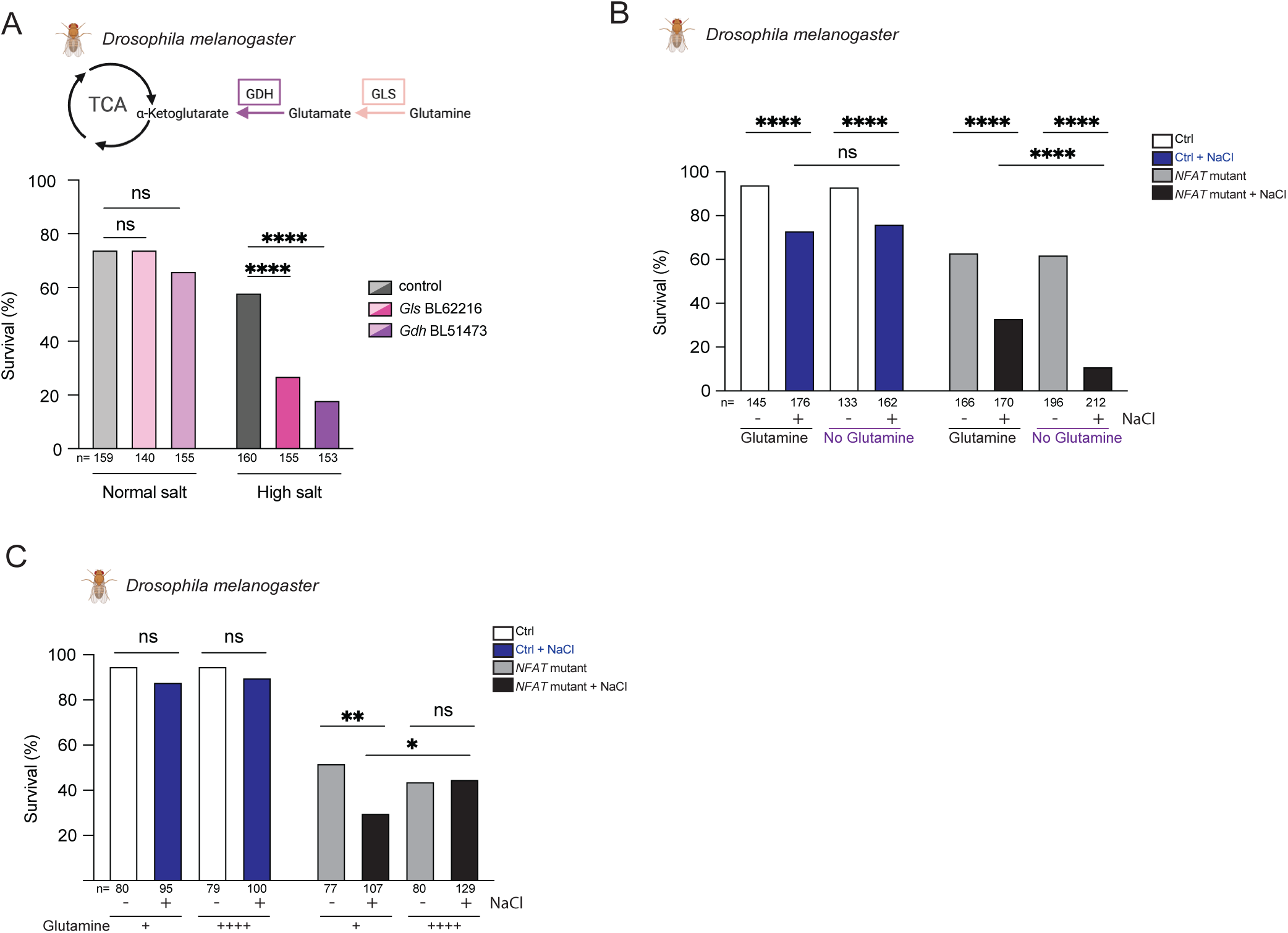
Glutamine is required for an appropriate response to hypertonic stress *in vivo*. (A) Survival analysis of D. melanogaster *Gls* (BL62216) and *Gdh* (BL51473) renal epithelial knockdown flies when exposed to NaCl hypertonic stress and water restriction. (B, C) Survival analysis of control or NFAT mutant D. melanogaster after indicated feeding paradigms (fig S8). Data in A,B, and C, presented as % survival analyzed with Fischer’s exact test, *p<0.05, ** p<0.01, *****p<0.0001. n as indicated in each panel.

To further characterize the interaction between hypertonic stress and glutamine metabolism *in vivo*, we examined *Drosophila melanogaster* mutants. *Drosophila* have a single *NFAT* gene that is most homologous to mammalian *NFAT5* ^34^. Flies with a large deletion in *Drosophila NFAT, NFAT^Δab^* ^34^, had higher lethality than controls under water-restricted conditions, which was worsened by hypertonic stress from high salt feeding and further exacerbated by dietary glutamine restriction (Fig 6B, fig. S8). In contrast, wet starvation conditions (i.e., access to water but no food) had no effect on mortality in any group, confirming that mortality is driven by lack of water rather than lack of calories (fig. S8, S9). Conversely, dietary supplementation with an additional 25 mM glutamine for seven days prior to water restriction abolished the excess lethality seen with high salt exposure in *NFAT* mutant flies (Fig. 6C), further highlighting the need for both glutamine and NFAT function in the response to hypertonic stress.

Together, these results show that hypertonic stress leads to lipid synthesis and accumulation across biological models, highlighting an evolutionarily conserved mechanism to increase potential water storage.

## Discussion

The ability to efficiently generate water from lipids during periods of water restriction provides a survival advantage. Desert animals, plants, and marine mammals have developed the ability to maximize the generation of water from lipids. However, despite association data between water restriction and lipid metabolism ^3, 8, 14, 27, 35, 36, 37, 38^, whether and how water-restriction-induced hypertonic stress prompts lipid accumulation was unknown. In this study we show that dehydration promotes lipid synthesis.

Our *in vitro* and *in vivo* data show the accumulation of lipids occurs when cells and animals are exposed to hypertonic stress. Our untargeted lipidomic analysis indicates that ether-linked long-chain triacylglycerols and cardiolipins accumulate under hypertonic stress and that the transcriptional response promotes lipid synthesis and elongation. Moreover, the enrichment of triglycerides with ether linkages at the sn1 position suggests a more stable form of storage that is resistant to hydrolysis. It is possible that in parallel to increased lipid synthesis, hypertonic stress and NFAT5 may also inhibit lipid breakdown (as previously shown in the heterozygous NFAT5 KO mice^28^). Thus the overall net effect being lipid accumulation from both increased synthesis and decreased breakdown.

We now provide a unifying explanation that links the decreases in OCR, changes to mitochondrial metabolism, and lipid accumulation seen during hypertonic stress. Our data is consistent with prior studies which saw decreased cellular OCR and pyruvate dehydrogenase inhibition after exposure to hypertonic stress ^27, 38, 39^, as well as others which had shown the accumulation of lipids in animals and plants exposed to hypertonic stress ^6, 7, 40, 41^. The inhibition of AMPK and activation of ACC which promote lipid synthesis in response to hypertonic stress suggest that despite a decreased OCR, cells can avoid an energy deficit that would otherwise blunt the lipogenic response. This would explain the changes we see in mitochondrial morphology, as mitochondrial elongation allows for more efficient energy production. Additionally, mitochondrial elongation can prevent mitochondrial degradation and promote cell survival during cellular stress^26^. Thus, our observations may represent an adaptive strategy by the mitochondria to maintain both efficient ATP production and cell viability.^42^

Our data also show that the dynamic metabolic profile of cells under hypertonic stress is regulated by the transcription factor NFAT5, which exerts pro-lipogenic effects. Prior work had shown that the partial absence of NFAT5 promoted increased oxygen consumption and prevented lipid accumulation in mice fed a high fat diet ^28^. We now build on these observations and demonstrate a role for NFAT5 as a critical regulator of fat metabolism in response to hypertonic stress. Whether there is an additive effect in which high-fat feeding and hypertonic stress can synergistically increase NFAT5-mediated obesity is currently unknown.

We found that glutamine incorporation into the TCA was required to synthesize lipids in response to hypertonic stress. Cancer cells are known to utilize glutamine for lipid synthesis^32, 33^. However, in contrast to cancer cells that reductively carboxylate glutamine, we found that glutamine is oxidized during hypertonic stress. However it still unclear how glutamine oxidation is maintained despite decreases in oxygen consumption rate. There is prior data that suggests glutamine can maintain TCA cycling and promote lipid synthesis when pyruvate transport is inhibited, akin to what we see during hypertonic stress in cells ^43^. Furthermore, our *Drosophila melanogaster* data highlights the critical role that glutamine plays in the response to hypertonic stress *in vivo*.

Prior work showed that when water restricted, desert adapted mice (*Notomys alexis*), shift to beta-oxidative metabolism, while BALB/c laboratory mice synthesize and accumulate fat over several days ^41^. We now show that within 24 hrs of water restriction, C57BL6/J mice increase fat mass similar to what was observed in BALB/c mice.Allen et al. showed that chronic dehydration remodels metabolism and increases metabolic water production in mice, however the time line in which animals can switch between predominantly lipid-synthesis or beta-oxidative metabolism is unclear ^44^.Thus, the duration of the dehydration stress on lipid synthesis and the timing of lipid synthesis and breakdown remain key areas to explore. In summary, our data now provide a direct link between dehydration, hypertonic stress, and lipid accumulation.

## Acknowledgements

We thank T. Blackwell, A. Morrison, and J. Gavin for the critical reading of the manuscript. *Drosophila* stocks obtained from the Bloomington *Drosophila* Stock Center (NIH P40OD018537) were used in this study. Additional stocks were obtained from the Vienna *Drosophila* Resource Center, part of the Vienna Biocenter Core Facilities, a non-profit research infrastructure publicly funded by the Austrian Federal Ministry of Education, Science and Research, and the City of Vienna via the Vienna Business Agency. We are grateful to Drs. Carl Thummel, Julian Dow, Shireen Davies, and Adrian Rothenfluh for sharing *Drosophila* fly lines used in this study.

## Funding

National Institutes of Health grant K08-DK135931 (to JPA)

National Institutes of Health grant R01-DK110358, R01-DK098145 (to ARR)

National Institutes of Health grant AR01DK105550 (to JFR),

National Institutes of Health grant DP5-OD033412 (to AST),

National Institutes of Health grant K08-DK134879 (to FB),

National Institutes of Health grant R38-HL167237 (to JSC),

National Institutes of Health grant DK51265, DK95785, DK62794, (to RCH and MZ),

National Institutes of Health grant DK069921 and DK127589 (to RZ);],

Veterans Affairs Merit Award 00507969 (to RCH),

Veterans Affairs Merit Award I01-BX002196 (to RZ),

National Institutes of Health grant R01-DK081646 (to VHH),

Intramural Research Program of the NIH, NIEHS ES103361-01 (to JAW),

National Institutes of Health grant R21-DK137147 (to JDY),

National Institutes of Health grant R35GM146951 (to JPVM),

National Institutes of Health grant K01-DK135924 (to CMH)

National Institutes of Health grant T32CA009582 (to EQJ),

National Institutes of Health grant DK135073, DK020593 (to LL),

National Institutes of Health grant 5K08DK133496 (to IC),

Veterans Affairs Merit Award EB033676, BX004258 (to MHW).

Hakim Family Foundation Physician Scientist Development Fund (JPA, AST, FMB)

Winkler Family Foundation (JPVM)

Destination Biochemistry Postdoctoral Scholars Program (EQJ)

Department of Defense National Defense Science and Engineering Graduate Fellowship Program (ERP)

Breakthrough T1D Advanced Postdoctoral Fellowship (KV)

Canadian Institutes of Health Research PJT 180476 (PK)

Agriculture and Food Research Initiative Foundational Knowledge of Plant Products (A1103) program, U.S. Department of Agriculture’s National Institute of Food and Agriculture, project award no. 2024-67014-42517 and 2021-67013-34009 (D.K.K.)

Fellowship grant from the Uehara Memorial Foundation (RB)

Sharon Anderson fellowship award from the American Society of Nephrology (RB)

Robert Wood Johnson Foundation Harold Amos Medical Faculty Development Program (JPA)

Research Scholar Award from the Southern Society for Clinical Investigation (FB)

The Digital Histology Shared Resource core at Vanderbilt University Medical Center (https://www.vumc.org/dhsr), the Translational Pathology Shared Resource core (P30-CA68485), and the Shared Instrumentation grant S10-OD023475. This work is supported by Metabolomics Workbench/National Metabolomics Data Repository (NMDR) (grant# U2C-DK119886), Common Fund Data Ecosystem (CFDE) (grant# 3OT2OD030544) and Metabolomics Consortium Coordinating Center (M3C) (grant# 1U2C-DK119889). This work was supported in part using the resources of the Center for Innovative Technology (CIT) at Vanderbilt University.

Body composition and pair-feeding experiments were carried out in the Vanderbilt Mouse Metabolic Phenotyping Center (DK135073, DK020593).

Financial support for this work provided by the NIDDK Innovative Science Accelerator Program (ISAC, www.isac-kuh.org), grant DK128851 (JPA)

## Author contributions

**Conceptualization**: JPA, ARR, JCR, FB, AST, JAW, JSC

**Investigation**: JSC, JPA, ARR, JS, YZ, HJN, CP, GN, EQJ, KV, ETA, JTT, WSY, PK, MM, OV, ERP, JPVM, RB, VHH, SC, KLL, SGC, JBT, CMH, LL, DKK, MW, DS, DAD, CMH, LL, MHW

**Methodology**: JSC, JPA, ARR, EQJ, JTT, JPC, MAC, JAW

**Formal Analysis**: JPA, JSC, ARR, FB, EQJ, KV, WSY, ASM, JPC, MAC, KLL, ACS, JAW, YZ, PK, IC, DKK

**Resources**: JPA, ARR, MHW, RCH, MZ, SDS, JAM, PK, JAW

**Project administration**: JPA, ARR, JCR, SDS

**Visualization**: JPA, ARR, FB, ASM, KLL, SGC, HJN, CP, WSY, PK, DKK

**Writing – original draft**: JPA

**Writing – review and editing**: JPA, ARR, JCR, JPC, EQJ, KV, ASM, JDY, KLL, SGC, ACS, SDS, JAM, RZ, RCH, JAW, AST, FB, IC

## Competing interests

Authors have no competing interests.

## Data and materials availability

The untargeted lipidomics data is available at the NIH Common Fund’s National Metabolomics Data Repository (NMDR) website, the Metabolomics Workbench, https://www.metabolomicsworkbench.org, where it has been assigned Project ID PR002443. The data can be accessed directly via its Project DOI: http://dx.doi.org/10.21228/M8VG1V.

RNA sequencing data is available at GSE274371.

CUT and Tag data is available at GSE302740. All other data are available in the manuscript or the supplementary materials.

## Notes

### Competing Interest Statement

The authors have declared no competing interest.

### Summary of Updates

New Figure 2, panel 4C, supplemental files updated and discussion materials related to peer review suggestions.

