## Supplemental Material for "Dehydration promotes intracellular lipid synthesis and accumulation"

**The PDF file includes:**

Materials and Methods
Supplementary Text
Figs. S1 to S10
Tables S1

### Materials and Methods

#### Cell culture

##### Kidney collecting duct cells

Inner Medullary Kidney Collecting Duct Cells (CDs) have been reported previously [45, 46]. Briefly, WT C57Bl/6J kidney collecting duct cells were isolated and transfected with SV40 (large T-cell antigen). Prior to experiments cells were cultured in DMEM with low glucose, glutamine, and pyruvate (Gibco 11885), supplemented with 10% FBS, 100 IU/mL penicillin - 100 mg/mL streptomycin (Gibco 15140122), and Normocin 100 mg/mL (InvivoGen) and grown to 90% confluency. Cells from passages 2-10 were used for all experiments.

In vitro NaCl hypertonic stress - For all experiments requiring NaCl hypertonicity stress, twelve-hours prior to lysis, IMCD medium was changed to FBS and antibiotic/antimycotic free low glucose DMEM with an added 100 mmol NaCl or water to preserve nutrient dilution e.g., 900 mL of DMEM and 100 mL of either water or 1 M NaCl. Cells were kept in this medium for 12 hours followed by downstream experiments. Medium glucose was measured with the Assure Platinum Glucose meter (Arkray), medium glutamine was measured with a Glutamine Assay kit (Sigma Aldrich MAK438) according to the manufacturers instructions.

Immunoblotting **-** Protein was extracted from cells using RIPA buffer (Thermo Fisher Scientific) with protease and phosphatase inhibitors (Roche). Total protein was then quantified with BCA Assay (Pierce). Equal amounts of protein were loaded in MiniProteanTGX polyacrylamide precast gels (Bio-Rad) and transferred to nitrocellulose using Transblot Turbo (Bio-Rad). Nitrocellulose membranes were stained for total protein using Ponceau-S (Thermo Fisher Scientific) and blocked with 5% nonfat dry milk for 1 hour at room temperature. Primary and secondary antibodies used are listed in Supplemental Table 2. Membranes were developed with Immobilon ECL Ultra Western HRP Substrate (Merck Millipore) and imaged in an iBright 1500 (Thermo Fisher Scientific). Band density quantification was performed with ImageJ (NIH).

siRNA transfection - IMCDs were plated at a density $7.5 \times 10^5$ cells per well. Upon reaching 30% confluency, they were transfected with either Silencer Select (Thermo Fisher Scientific) *siNFAT5* (AssayID 73277), or ctrl siRNA (AssayID 4390843) according to the manufacturer's instructions for 36-48 hrs. Cells were treated with 100 mmol NaCl or control for 12hrs after which cells were used for downstream analysis i.e., cell imaging, metabolic profiling, or immunoblotting.

Imaging - Cells were seeded at a density of 100,000 cells/mL in 300 $\mu$L per well in an 8-well $\mu$-Slide glass slide (Ibidi) with supplemented DMEM growth media and allowed to grow for over 24 hours until reaching 90% confluency. Cells were then treated with serum/drug free DMEM with or without 100 mmol added NaCl for 12 hrs. For visualization of lipids either BODIPY or oil-red-o were added according to the manufacturer's instructions. Briefly, BODIPY (Invitrogen D3922) was added to each well according to manufacturer's instructions and incubated for 30 minutes prior to imaging. For oil-red-o (Abcam ab150678), after incubation with NaCl or control medium, cells were washed with PBS and fixed with 4% PFA for 30 min at room temperature followed by incubation with oil-red-o. For mitochondria staining, the MitoTracker probe (Cell Signaling 9074P) or MitoSox Red (Invitrogen M36008) were used according to the manufacturer's instructions. Briefly, prior to microscopy, a 1 mM stock solution of MitoTracker Green FM (50 $\mu$g lyophilized solid reconstituted in 74.4 $\mu$L DMSO) was diluted into phenol free DMEM media to a final concentration of 250 nM which was then added to cells. For MitoSox

Red, a 5mM stock solution of MSR was diluted to create a 500 nM working solution which was then added to the cells. In both cases, cells were incubated at 37°C for just under 30 minutes prior to imaging.

Cells were imaged with a Nikon TiE fully motorized inverted fluorescent microscope or a Zeiss LSM 980 Confocal AiryScan 2. For BODIPY staining and MitoTracker excitation/emission were 493/504 and 490/516 respectively. For oil-red-o slides were scanned with the Aperio VERSA 200 pathology scanning system (Leica Biosystems).

Image Analysis - Oil-red-O particle staining was performed with ImageJ. Aperio scans (400 uM x 400 uM) were analyzed with particle analysis post manual threshold to red to create binary images. Three images were analyzed per biologic replicate from four independent experiments.

BODIPY analysis for both CDs and elephant seal cells was carried out with NIS elements software. A square region of interest (ROI) of $200\mu m^2$ was selected and manually thresholded, at least 10 ROIs were taken per biologic replicate from three independent experiments. BODIPY intensity was normalized to the control image from each independent experiment.

For mitochondrial aspect ratio analysis, we used the “Mitochondria Analyzer” plugin for ImageJ/Fiji as previously described [47]. Two to three images were analyzed from 3 independent experiments.

Oil-Red-O staining – whole well absorbance – When BODIPY staining quantification was deemed difficult due to inability to quantify individual cells in a non biased manner, we used whole well cell number adjusted Oil-Red-O absorbance at 492 nm to quantify total lipids in treated cells. Therefore, IMCD cells were seeded into a 24-well plate in culture media at a density of 10,000 cells per well in 1 mL of cell suspension and cultured for 72 hours. Subsequently, cells were treated with or without glutamine or *siNADK2* and siCtrl and 100 mM NaCl in media without FBS or antibiotics/antimycotics for 12 hours prior to Oil-Red-O staining. On the day of staining, an aliquot of Oil-Red-O stain was pre-heated to 60°C. Cells were fixed with 4% PFA for 15 minutes, 1 mL 60% IPA was added, and the plate was incubated for 5 minutes at room temperature. All wells were aspirated, and cells were allowed to dry completely at room temperature. 500 μL of pre heated and filtered Oil-Red-O stain was added to each well and incubated at room temperature for 15 minutes. The stain was removed, and each well was washed 3 times with 1 mL water until clear. Water was aspirated, cells were allowed to dry completely, stain was eluted with 250 μL IPA per well, and the plate was incubated for 10 minutes at room temperature on a rocker. 200 μL of the solution was transferred to a 96-well plate, and absorbance was read at 492 nm, using 200 μL of 100% IPA as a blank. Data were normalized to cell number as determined using the automated BioRad TC-20 cell counter.

Metabolic flux analysis - IMCD cells were seeded into microplates at a density of 20,000 cells/well. Mito Stress Test kits (Agilent Technologies) were used to measure OCR with the Agilent Seahorse XFe24 metabolic flux analyzer, according to the manufacturer’s instructions. Data were normalized to cell number as determined using the automated BioRad TC-20 cell counter.

Stable Isotope Labeling (SIL) Targeted Metabolomics [48] - IMCD cells were plated in 10 $cm^2$ culture dishes in serum free, glutamine free DMEM and 4.05 mM $^{13}C_5$-glutamine (Cambridge Isotope Laboratories, Tewksbury, MA) with +/- 100 mmol NaCl for 12 hrs. After treatment with heavy labeled glutamine, media was removed, and wells were washed twice with room temperature PBS. After washing, cells were scraped from the plate in 3 mL PBS, and transferred to a 15 mL conical tube. The cells were then pelleted via centrifugation and PBS was removed. Samples were then flash frozen in liquid nitrogen and stored at -80°C. To extract metabolites, 1 mL of -80°C 80:20 MeOH:$H_2O$ was added to the cells and samples were vortexed vigorously. 100 nmol $^{13}$C-1-Lactate (internal standard, MilliporeSigma, Burlington, MA) was then spiked into each sample. Cells were then placed in a -80°C freezer to extract for >15 minutes. Precipitated protein from extracted cells was pelleted at 2,900 x *g* for 10 minutes at 4°C. Supernatants containing metabolites were then transferred to new 15 mL conical tubes and dried under nitrogen. Precipitated protein pellets were resolubilized and protein concentration was measured via BCA assay (Thermo Fisher Scientific, Waltham, MA). Dried samples were resuspended in 50 $\mu$L 3:2 mobile buffer A: mobile buffer B (see below). 18 $\mu$L of the sample was then chromatographed with a Shimadzu LC-30 system equipped with a 100 x 2.1mm, 3.5µm particle diameter XBridge Amide column (Waters, Milford, MA). Mobile phase A: 20 mM $NH_4OAc$, 20 mM $NH_4OH$, 5% acetonitrile in $H_2O$, pH 9.45 (pH adjusted with $NH_4OH$); mobile phase B: 100% acetonitrile. With a flow rate of 0.45 mL/min the following gradient was used: 2.0 min, 95% B; 3.0 min, 85% B; 5.0 min, 85% B; 6.0 min, 80% B; 8.0 min, 80% B; 9.0 min, 75% B; 10 min, 75% B; 11 min, 70% B; A12 min, 70% B; 13 min, 50% B; 15 min, 50% B; 16 min 0% B; 17.5 min, 0% B; 18 min, 95% B. The column was equilibrated for 3 minutes at 95% B between each sample. Scheduled MRM was conducted in negative mode with a detection window of 120 seconds using an AB SCIEX 6500 QTRAP with the following analyte parameters: *m/z* 191 → 111 for citrate/isocitrate (retention time (RT) 13.4 minutes); *m/z* 193 → 113 for $^{13}C_2$-citrate/isocitrate (RT 13.4 minutes); *m/z* 195 → 114 for $^{13}C_4$-citrate/isocitrate (RT 13.4 minutes); *m/z* 196 → 116 for $^{13}C_5$-citrate/isocitrate (RT 13.4 minutes); *m/z* 197 → 116 for $^{13}C_6$-citrate/isocitrate (RT 13.4 minutes); and *m/z* 90 → 43 for $^{13}$C-1-lactate [internal standard] (RT 5.5 minutes). All analytes were quantified via LC-MS/MS using the $^{13}$C-1-Lactate internal standard and normalized to the protein in each respective sample's cell pellet. Outliers were removed using interquartile range.

<u>CUT&Tag</u> – Kidney collecting duct cells grown in isotonic media (as above) and collected 1 hr after supplementation with 100 mM NaCL were harvested using StemPro Accutase Cell Dissociation Reagent (Thermofisher) and washed with PBS. Cells were counted and HK-2 (ATCC) cells were spiked in at a ratio of 1:100. CUT&Tag was performed as described previously [49]. Cells were washed with wash 150 buffer (final concentrations: 20 mM HEPES ($Na^+$) pH 7.5, 150 mM NaCl, 0.5 mM spermidine, 1X EDTA-free complete protease inhibitor) and added to BioMag Plus Concanavalin A Beads (Bangs Laboratories) pre-activated with cold bead activation buffer (20mM HEPES pH7.5, 10mM KCl, 1mM $CaCl_2$, 1mM $MnCl_2$) for 10 minutes on a rotator at room temperature. Wash buffer was removed and cold antibody buffer (final concentrations: Wash150 buffer, 100x Protease Inhibitor, 0.05% Digitonin (Sigma), 2 mM EDTA, 1% BSA (Sigma) was added. 1 $\mu$g of NFAT5 (Bethyl, Rb pAb, A305-174A), H3K27ac (Active motif, Ms mAb, 39685), or H3K27me3 (Active motif, Rb pAb, 39155) antibodies were added to each reaction and incubated overnight with orbital mixing at 4°C. Reactions were washed with Digitonin150 buffer (final concentrations: Wash150 buffer, 100xPI, 0.05% digitonin) three times. 1 $\mu$g of secondary antibody Guinea pig polyclonal anti-rabbit IgG (ABIN101961) or Rabbit Anti-Mouse IgG H&L (ab46540) was added and incubated on an orbital rotator at room temperature for 1 hr. Reactions were washed with Digitonin150 buffer three times. 2.5 $\mu$l of pAG-Tn5 fusion protein (Epicypher) was diluted with Digitonin300 buffer (20 mM HEPES pH 7.5, 300 mM NaCl, 0.5 mM spermidine, 100x Protease inhibitor, 0.01% Digitonin), then added to each reaction and incubated for 60 minutes at room temperature. After reaction was washed with Digitonin300 buffer three times, tagmentation buffer (Digitonin300 buffer, 10 mM $MgCl_2$) was added to reactions and incubated at 37°C for 1 hr. To stop tagmentation, buffer containing 4.2 $\mu$l 0.5 M EDTA, 1.25 $\mu$l 10%SDS, 1.1$\mu$l Proteinase K (20mg/mL) was added to the reaction and incubated for 1 hr at 55°C. DNA was purified using a MinElute PCR Purification Kit (Qiagen) as per manufacturer's instructions.

The product was PCR amplified using barcoded primers (See supplementary table) and NEBNext HiFi 2x PCR master mix (NEB). After PCR, libraries were purified using AMPure XP (Beckman) at 1:1 ratio and 80% ethanol. Libraries were eluted in DNase/RNase free $H_2O$. Libraries were quantified using a Qubit fluorometer and dsDNA HS assay kit (Thermofisher) and then analyzed using Agilent Tapestation, and sequenced on a Nextera sequencer (Illumina).

Paired-end fastq files were aligned to the mouse mm10 reference genome with bowtie2 version 2.5.2 [50] after removing low quality reads with fastp [51]. The alignment parameters were (bowtie2 -p 8 -X 700 --local --very-sensitive-local --no-unal --no-mixed --no-discordant --phred33 -I 10). Aligned .sam files were converted to coordinate sorted binary .bam files and duplicates removed with picard SortSam and MarkDuplicates version 2.25, respectively, https://broadinstitute.github.io/picard/. Normalized bigWig read coverage files were created using the deepTools bamCoverage application [52]. Scale factors for normalization were determined using hg38 reads obtained from spiked-in HK2 human kidney cells for each sample. Data are plotted as reads per genomic content (RPGC). To quantify enrichment at specific promoter sites, read coverage was determined near each gene TSS and centered at the NFAT5 peak summit. Reads were obtained for histone marks H3K27ac and H3K27me3 at the same sites. Read coverage plots were created inR using “rtracklayer”, “GenomicRanges”, “BSgenome.Mmusculus.UCSC.mm10” and “ggplot2” packages. Data is available at GSE302740.

Bulk RNA sequencing - IMCDs were treated with 100 mmol NaCl or control medium for 12 hrs. RNA was then extracted with Trizol (Invitrogen). A Quality Control analysis was performed on the RNA samples using the Agilent Bioanalyzer to assess quality and an RNA Qubit assay was used to determine quantity. Poly(A) RNA enrichment was conducted using NEBNext Poly(A) mRNA Magnetic Isolation Module (NEB), and the sequencing library was constructed by using the NEBNext Ultra II RNA Library Prep Kit (NEB) (P/N: E7765L, NEB, Ipswich, MA) following the manufacturer’s instructions. End repair, A-tailing, and adapter ligation was performed to generate the final cDNA library. The library quality was assessed using a Bioanalyzer and quantified using a qPCR-based method with the KAPA Library Quantification Kit (P/N: KK4873) and the QuantStudio 12K instrument. Prepared libraries were pooled in equimolar ratios, and the resulting pool was subjected to cluster generation using the NovaSeq 6000 System, following the manufacturer's protocols. 150 bp paired-end sequencing was performed on the NovaSeq 6000 platform targeting 50M reads per sample. Raw sequencing data (FASTQ files) obtained from the NovaSeq 6000 was subjected to quality control analysis, including read quality assessment. Real Time Analysis Software (RTA) and NovaSeq Control Software (NCS) (1.8.0; Illumina) were used for base calling. MultiQC (v1.7; Illumina) was used for data quality assessments.
Paired-end RNA sequencing reads (150bp long) were trimmed and filtered for quality using Trimgalore v0.6.7 [53]. Trimmed reads were aligned and counted using Spliced Transcripts Alignment to a Reference (STAR)[54] v2.7.9a with the –quantMode GeneCounts parameter against the GRCm38 (mm10) mouse genome. DESeq2 package v1.36.0 [55] was used to perform sample-level quality control, low count filtering, normalization and downstream differential gene expression analysis. Genomic features counted fewer than five times across at least three samples were removed. False discovery rate adjusted for multiple hypothesis testing with Benjamini-Hochberg (BH) procedure $p$ value $< 0.05$ and log2 fold change >1 was used to define differentially expressed genes. Pairwise comparison with NaCl-treated cells compared to non-treated cells was performed for differential analysis. Four biological replicates per condition were included for the differential expression analysis. A ranked list of

these results was used as input for Gene Set Enrichment Analysis using ClusterProfiler [56] and KEGG metabolic pathways [57] were filtered with the "mmu00" prefix. Data is available at GSE274371.

*Untargeted Lipidomics.*

Mass Spectrometry Lipidomic Standards**.** Standards for mass spectrometry analysis were used for quality assurance and control (to assess sample preparation variability and instrument variability) were procured from Avanti Polar Lipids (Birmingham, AL) and Cayman Chemical (Ann Arbor, MI). SPLASH Lipidomix Mass Spec Standard (Avanti Polar Lipids) was used to assess sample preparation and processing variability. Standards to assess instrument variability included four exogenous lipid standards: N-heptadecanoyl-D-erythro-sphingosine (C17 ceramide (d18:1/17:0), Avanti Polar Lipids), D-glucosyl-b1-1'-D-erythro-sphingosine (glucosyl(b) sphingosine (d18:1), Avanti Polar Lipids), heptadecanoic acid (FA 17:0, Cayman Chemical), and nonadecanoic acid (FA 19:0, Cayman Chemical).

Lipidomics Sample Preparation and Lipid Extraction**.** Frozen samples were thawed on ice and lysed in ice-cold lysis buffer (1:1:2, acetonitrile: methanol: ammonium bicarbonate 0.1M, pH 8.0) to an equal cell density per sample, followed by probe tip sonication with 10 pulses at 30% power. Samples were normalized by total protein amount (150$\mu$g) based on a bicinchoninic acid (BCA) assay and further mixed with 800$\mu$L of cold methanol, vortexed for 30 seconds and incubated overnight at -80°C for protein precipitation. Following incubation, samples were centrifuged for 10 minutes at 15,000 rpm at 4°C and the supernatant was transferred and dried down using a vacuum centrifuge.

Samples were reconstituted in 100 $\mu$L $H_2O$, 100 $\mu$L methanol, and 10 $\mu$L of SPLASH Lipidomix with vortex mixing after each addition. Samples were incubated at room temperature for 10 minutes followed by liquid-liquid extraction (LLE). For LLE, 600 $\mu$L MTBE was added with vortex mixing for 30 seconds followed by incubation on ice for 10 minutes and centrifugation at 15,000 rpm for 15 minutes at 4°C. The upper (hydrophobic) fraction was transferred and dried down using vacuum centrifugation and stored at -80°C for further lipidomic studies. Prior to mass spectrometry analysis, individual hydrophobic extracts were reconstituted in 8 0$\mu$L methanol:chloroform (9:1, v:v) containing exogenous lipid standards. A pooled quality control (pooled QC) sample was prepared by pooling equal volumes (30 µL) from each individual sample following reconstitution.

Following processing and normalization, untargeted lipidomic data were analyzed within LipidSuite[58] Data presented are from positive ion mode. Class enrichment reported after multiple comparisons correction. Significance thresholds were set at 0.05

Liquid Chromatography-Mass Spectrometry Data Acquisition for Untargeted Lipidomics**.** Prepared samples were analyzed by reversed-phase liquid chromatography-tandem mass spectrometry (RPLC-MS/MS) in the Center for Innovative Technology (CIT) at Vanderbilt University using previously described methods [59]. Briefly, chromatographic separations were performed with an Agilent 1290 Infinity LC system with a Hypersil Gold column (1.9 $\mu$m, 2.1 mm x 100 mm) at 40°C. The flow rate was kept at 0.25 mL/min across the LC gradient with an injection volume of 5 µL. Mobile phase A was $H_2O$, mobile phase B was 60:36:4 (v:v:v) acetonitrile:isopropanol:$H_2O$, and both mobile phases had additives of 10 mM $NH_4OAc$ and 0.1% formic acid. The Agilent 6560 instrument was fitted with electrospray ionization (Dual

AJS, Agilent) and source conditions were optimized for lipidomic analyses and operated in both positive and negative ionization modes with gas temperature of 280°C, drying gas flow of 5 L/min, nebulizer at 10 psi, sheath gas temperature of 300°C, sheath gas flow at 11.8 L/min, capillary at 3500 V, nozzle at 2000 V, and octupole RF at 750 $V_{pp}$.

The Agilent 6560 instrument was tuned in low (1700 *m/z*) mass range with a prepared 10% solution of ESI-L Low Concentration Tuning Mix (Agilent). Solvent blanks and standard blanks were acquired at the beginning of the analysis to assess spectral contaminants and to generate an exclusion list for the MS/MS method. The pooled QC sample was injected five times immediately preceding the batch to condition the LC column, and again after every five samples to assess instrument repeatability through principal component analysis. Samples were injected in randomized order, and 10% of samples were reinjected in each polarity for quality control assessment. Fragmentation spectra were acquired in each polarity via three iterative, top two, data dependent analysis MS/MS acquisitions of the pooled QC sample.

Lipidomics Data Analysis**.** Mass spectrometry data was imported, processed, and normalized to all compounds using Progenesis QI v.3.0 (Non-linear Dynamics, Newcastle, UK). All MS and MS/MS sample runs were aligned against a pooled QC sample. QA/QC metrics were assessed throughout the experiment by monitoring retention times and peak areas of SPLASH Lipidomix standards. Relative abundances for both positive and negative ion mode data acquisitions were less than 14%CV for 90% of standards for sample preparation and less than 5%CV for 100% of standards for instrument variability. Annotations [60] were assigned using accurate mass measurements (<10 ppm), isotope distribution similarity scores, and assessment of MS/MS spectrum matching (when applicable) from MS Dial database searching, as well as MS/MS searches to Lipid Annotator [61] and Lipid Match [62] Lipid species significance was assessed by ANOVA (analysis of variance) from normalized compound abundance data. Visualizations of lipidomics annotations and statistics were prepared using Cytoscape 3.10.3 [63]. Global visualizations including Principal Component Analysis (PCA) were created using MetaboAnalyst v6.0 [64] using Pareto scaled, log-transformed data.

Data Availability - The untargeted lipidomics data is available at the NIH Common Fund's National Metabolomics Data Repository (NMDR) website, the Metabolomics Workbench, https://www.metabolomicsworkbench.org, where it has been assigned Project ID PR002443. The data can be accessed directly via its Project DOI: http://dx.doi.org/10.21228/M8VG1V.

**Elephant seal endothelial cells**

Northern elephant seal primary vascular endothelial cells were derived and characterized following our previously published methods [65] under NMFS permits 19108 and 22479. Cells were grown in low-glucose DMEM (Gibco) supplemented with 10% fetal bovine serum (Seradigm), 10 mM HEPES (Gibco), 1% Antibiotic–Antimycotic (Gibco cat), and 4 µg/mL endothelial cell growth supplement (Corning). Cells seeded into 6-well plates were incubated with 100 mM NaCl in serum-free medium for 12 hours (21% O2, 37°C, 5% CO2) and lysed with Pierce RIPA Buffer (Thermo Scientific) containing 2% Halt Protease and Phosphatase Inhibitor Cocktail (Thermo Scientific cat # 78440). Protein content was measured using a Pierce BCA Rapid Gold Protein Assay (Thermo Scientific cat # A55862). Proteins were separated using 4-12% Bolt Bis-Tris Plus gels (Thermo Scientific cat # NW04120BOX) and transferred onto nitrocellulose membranes. For imaging, BODIPY (Invitrogen D3922) was added to each well according to manufacturer's instructions and incubated for 30 minutes prior to imaging. Samples were visualized using a Zeiss Axio Observer 7 inverted microscope fitted with a ×63 objective and Zen software (Zeiss, Oberkochen, Germany).

**HeLa Cells**

*Cell culture*

HeLa cells were obtained from ATCC (Ref. No. CCL-2) and cultured in DMEM (Gibco, 11885-084) supplemented with 5% FBS. For serum starvation, DMEM without FBS supplementation was used.

*Live imaging*

Cells were grown in Lab-Tek™ II chambered coverglass (Nunc). Live imaging was performed using the Zeiss LSM980 laser-scanning confocal microscope with the appropriate lasers and filters. For general intensity measurement, a 20×0.8 NA Plan-Apochromat objective was used. To acquire super-resolution images for morphological analysis, a 63×1.4 NA Oil Plan-Apochromat objective was used with AiryScan 2. The images were processed and deconvoluted using the built-in AiryScan algorithms.

Cells were incubated with 100 nM MitoTracker, 100 nM TMRM, and/or 1 $\mu$g/mL BODIPY for 30 minutes to visualize mitochondria, mitochondrial membrane potential, and lipid droplets, respectively. Cells were washed twice with their respective media before imaging. Cells were treated with 10 $\mu$M CCCP for 10 minutes as a control for decreased mitochondrial membrane potential if indicated.

*Image analysis*

All images were analyzed using Fiji (ImageJ). In brief, to analyze the mitochondrial morphology, a maximum projection image was generated from the Z-stack images of fluorescently labeled mitochondria. A square region of interest (ROI) of 225 $\mu m^2$ was selected and manually thresholded. The mean area and number of mitochondria within the ROI were measured using the "Analyze Particles" plugin while excluding particles of <0.1 $\mu m^2$.

To measure the relative mitochondrial membrane potential ($\Delta\Psi$), a maximum projection image was generated from the confocal Z-stack images of TMRM- and MitoTracker-stained cells. Each cell is specified as the ROI in which the integrated density (total intensity) of the TMRM and MitoTracker signals were measured, respectively. The relative membrane potential in each cell is calculated by normalizing the TMRM signal to MitoTracker:

$$Relative\ \Delta\Psi = \frac{TMRM}{Mitotracker}$$

***Drosophila melanogaster***

*Drosophila melanogaster* strains used

Flies were raised on commercial glucose food (Archon Scientific, D20301) at 25°C unless otherwise indicated on a 12:12 hour light:dark cycle. *Drosophila melanogaster* strains obtained from the Bloomington *Drosophila* Stock Center (BDSC) and Vienna *Drosophila* Resource Center (VDRC) are listed below. BDSC 36304 was used as a control for experiments with BDSC 62216 and BDSC 51473. BDSC 36303 was used as a control for experiments with BDSC 51422. VDRC 60100 was used as a control for VDRC 108317, VDRC 109499, and VDRC 107032. Controls reflect the host strains containing the landing sites used to generate the RNAi lines. *w*Berlin (from Adrian Rothenfluh, University of Utah) was used a control for BDSC 32652. *w;* c42-*GAL4* [66] was obtained from Julian Dow and Shireen Davies (University of Glasgow) and combined with tubulin-Gal80$^{ts20}$ [67] using standard genetic techniques; both were outcrossed to *w*Berlin for 5 generations. For inducible adult Malpighian tubule knockdown of *Gls*, *Gdh,* and *NFAT*, *w;* tub-*GAL80$^{ts20}$*; c42-*GAL4* flies were crossed to RNAi lines listed below or the controls described above. Flies were reared at 18°C (GAL4 suppressed) throughout development and then transferred to 28°C (GAL4 activated) within 1-2 days of eclosion for 7 days to allow gene knockdown.

| BDSC stock number | Gene | Genotype |
|---|---|---|
| 36303 | n/a | $y^1$ $v^1$; P{CaryP}attP2 |
| 36304 | n/a | $y^1$ $v^1$; P{CaryP}$Msp300^{attP40}$ |
| 62216 | *Gls* | $y^1$ $sc^*$ $v^1$ $sev^{21}$; P{TRiP.HMC05223}attP40 |
| 51473 | *Gdh* | $y^1$ $v^1$; P{TRiP.HMC03217}attP40 |
| 51422 | *NFAT* | $y^1$ $sc^*$ $v^1$ $sev^{21}$; P{TRiP.HMC02411}attP2 |
| 32652 | *NFAT* | $w^{1118}$ $NFAT^{\Delta ab}$ |
| 7019 | n/a | $w^*$; P{tubP-$GAL80^{ts}$}20; TM2/TM6B, $Tb^1$ |

| VDRC stock number | Gene | Genotype |
|---|---|---|
| 60100 | n/a | y $w^{1118}$; P{attP $y^+$ $w^{3'}$} |
| 108317 | *Gls* | P{KK102885}VIE-260B |
| 109499 | *Gdh* | P{KK107890}VIE-260B |
| 107032 | *NFAT* | P{KK102385}VIE-260B |

For all Drosophila feeding paradigms see Supplementary figure 8.

Glutamine-deficient *Drosophila* diet

Chemically-defined food with (CDF) or without glutamine (glutamine-deficient diet, GDF) was prepared according to [68] and as listed in the table below. Fat mix was prepared fresh, typically in 20 mL batches. Sugar mix was made in 1L batches and stored at 4°C. 3-4 day old females, 10-15 per vial, were reared on CDF or GDF for 5 days at 25°C and then kept on CDF, CDF+0.3 M NaCl (Fisher, S271-3), GDF, or GDF+0.3 M NaCl for 24 hours followed by 24 hours in empty vials (Genesee Scientific, 32-116SB) or empty vials with water-saturated flugs (Genesee Scientific, 49-102). Living vs. dead flies were counted.

| Per liter | CDF | GDF |
|---|---|---|
| Essential Amino Acids (TD.10473) (g) | 11.50 | 11.50 |
| Non-essential Amino Acid Mix (TD.110036) (g) | 8.12 | |
| Non-essential Amino Acid Mix -Gln (TD.240605) (g) | | 6.92 |
| 5X Sugar Mix Solution (ml) | 160 | 160 |
| Sucrose (Fisher, S71204) – 398 (g/L) | | |

| | | |
|---|---|---|
| Lactose (Sigma, L2643) – 37 (g/L) | | |
| Glucose (VWR, BDH0230) – 30.7 (g/L) | | |
| Trehalose (Sigma, T9449) – 24.4 (g/L) | | |
| 100X Fat Mix Solution (ml) | 8 | 8 |
| Cholesterol (Sigma, C3045) - 0.199 (g/20mL) | | |
| Lecithin (MP Biomedicals,102147) – 1.98 (g/20mL) | | |
| Basal Mix (Vitamin & Mineral Mix; TD.130769) (g) | 1.70 | 1.70 |
| RNA (Sigma R6625) (g) | 1 | 1 |
| DNA (Sigma D6898) (g) | 0.5 | 0.5 |
| Agarose (Invitrogen 16500) (g) | 10 | 10 |

Glutamine-supplemented *Drosophila* diet

1-2 day old adult female flies were reared on Jazz mix *Drosophila* food (Fisher, AS153) with or without supplemental 25 mM L-glutamine (RPI, G36040) for 6-7 days and then flipped on to Jazz mix with or without supplemental 0.3 M NaCl and 25 mM glutamine, followed by 24 hours in empty vials with or without water-saturated flugs. Living vs. dead flies were counted.

Testing metabolic pathways required for *Drosophila* survival under osmotic stress/water restriction

8-9 day old adult female flies with inducible knockdown of metabolic enzymes were transferred to Jazz mix (Fisher, AS153) *Drosophila* food with or without 0.3 M NaCl supplementation for 24 hours at 28°C, with 10-15 females/vial. Flies were then flipped into empty vials with or without water-saturated flugs for 24 hours at 25°C. Living vs. dead flies were counted.

***Mus musculus***

Two mouse strains were used for this project, C57BL/6J mice obtained from Jackson laboratories and C57BL/6N-Atm1Brd Nfat5tm1a(KOMP)Wtsi/JMmucd, RRID:MMRRC_048793-UCD [69], which were obtained from the Mutant Mouse Resource and Research Center (MMRRC) at University of California at Davis, an NIH-funded strain repository, and was donated to the MMRRC by The KOMP Repository, University of California, Davis; Originating from Stephen Murray, The Jackson Laboratory.

*Water restriction.* Twenty-four hours prior to euthanasia, water was removed from cages of mice in the water-restricted group.
*Water loading.* Twenty-four hours prior to euthanasia, food in the water-loaded group was changed to a gelled diet. Briefly, 65 g crushed 4.5% fat mouse chow (LabDiets, 5L0D) was mixed with 7 g gelatin and dissolved in 120 mL water. Gel was solidified in plastic cups and then served as the sole source of food for 24 hours, 9:00 AM to 9:00 AM. Ad libitum access to water was maintained throughout.

*Pair-feeding* – Wild-type C57BL6/J mice were placed in separate metabolic cages and water was withheld. Food was weighed before and after to determine food consumption during water restriction, aprox. 3g per mouse per 24 hr period of water restriction. For the water loaded group, 3g of crushed 4.5% fat mouse chow was crushed and mixed with 1g of gelatin and 11 mL of water. Mice were then placed in metabolic cages and provided ad lib water and the gelled food. After water restriction or water loading body composition was determined by NMR (Bruker Minispec), mice were euthanized by cervical dislocation and blood collected for serum triglycerides (Cayman Item No. 10010303) and serum free fatty acids (Cayman Item No. 700310), which were both measured according to the manufacturers instruction.

*Osmolality* – Body fluid osmolality was measured using a freezing point osmometer from Precision Systems (No. 6002).

*Tissue lysate and serum triglyceride measurement* - Kidney lysate triglycerides were measured with the Triglyceride Colorimetric Assay Kit (Cayman Chemical Item No. 10010303) according to the manufacturer's instructions.

*Immunoblotting* **-** Protein was extracted from whole kidneys using RIPA buffer (Thermo Fisher Scientific) with protease and phosphatase inhibitors (Roche), total protein was then quantified with BCA Assay (Pierce), and equal amounts of protein were loaded in MiniProteanTGX polyacrylamide precast gels (Bio-Rad) and transferred to nitrocellulose using Transblot Turbo (Bio-Rad). Nitrocellulose membranes were stained for total protein using Ponceau-S (Thermo Fisher Scientific) and blocked with 5% nonfat dry milk for 1 hour at room temperature. Primary and secondary antibodies used are listed in Supplemental Table 1. Membranes were developed with Immobilon ECL Ultra Western HRP Substrate (Merck Millipore) and imaged in an iBright 1500 (Thermo Fisher Scientific). Band density quantification was performed with ImageJ (NIH).

**Statistics**

Data are reported as mean ± SD. Statistical analyses were performed with Prism 10 software (GraphPad Software Inc.), using a 2-tailed Student's t test, Mann-Whitney test, 1-way ANOVA with a post hoc Tukey's test, 2-way ANOVA with a post hoc Šidák's test, or Fischer's Exact Test, as indicated. P values of less than 0.05 were considered statistically significant. Statistical analyses for lipidomics and RNA sequencing are described above.

**Study approval**

All procedures involving mice were performed in accordance with the NIH Guide for the Care and Use of Laboratory Animals (National Academies Press, 2011) and were reviewed and approved by the Institutional Animal Care and Use Committee (IACUC) of Vanderbilt University, Nashville, Tennessee, USA.

Supplemental Figure 1

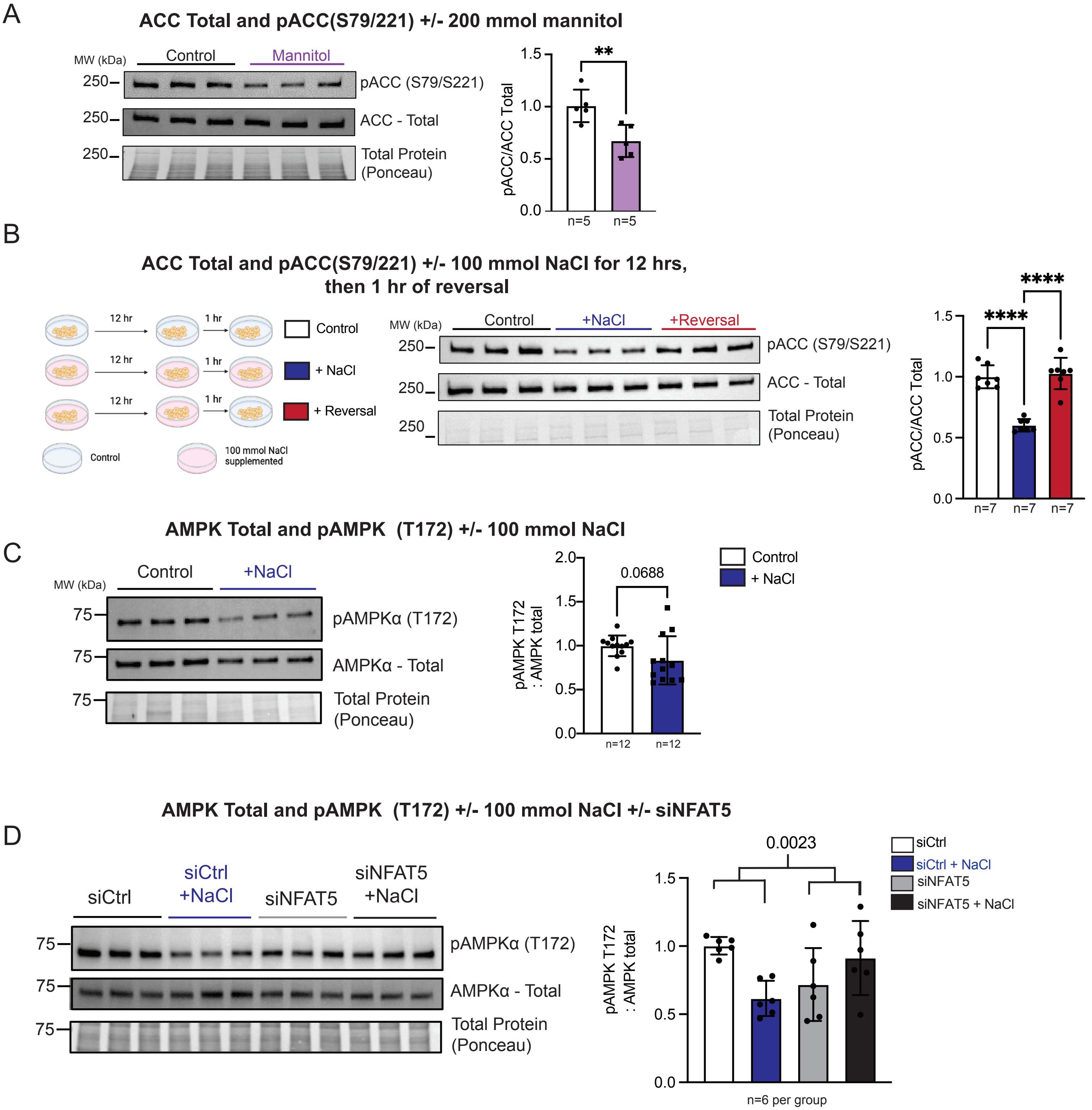


**Supplemental Figure 1. Hypertonic stress reduces phosphorylation of AMPK.** Immunoblots of total ACC and pACC (S79/221) from kidney collecting duct cell lysates treated with (A) 200 mmol mannitol vs. control for 12 hours or (B) 12 hours of 100 mmol NaCl followed by 1 hr of fresh 100 mmol NaCl medium (blue) vs 12 hours of 100 mmol NaCl followed by 1 hr of control medium (red) vs control (white). (C) Immunoblots of total AMP activated protein kinase (AMPK) and pAMPK (T172) from kidney collecting duct cell lysates treated with 100 mmol NaCl vs control. (D) Immunoblots of total AMP activated protein kinase (AMPK) and pAMPK (T172) in kidney collecting duct cells transfected with siNfat5 vs siCtrl treated with NaCl or control. Data in A, C presented as mean +/- SD, analyzed with a two-tailed Student's t-test, **p<0.01. n as indicated in each panel. Data in B presented as mean +/- SD, analyzed with One-way ANOVA with a post hoc Tukey test, **** p<0.0001. Data in D presented as mean +/- SD, analyzed with a two-way ANOVA with NaCl and siRNA as the independent variables. p=0.0023 for the interaction, siCtrl vs siCtrl +NaCl p=0.0075, siNFAT5 vs siNFAT5 + NaCl p=0.2152 after Šídák's multiple comparisons test. n as noted in figure.

Supplemental Figure 2

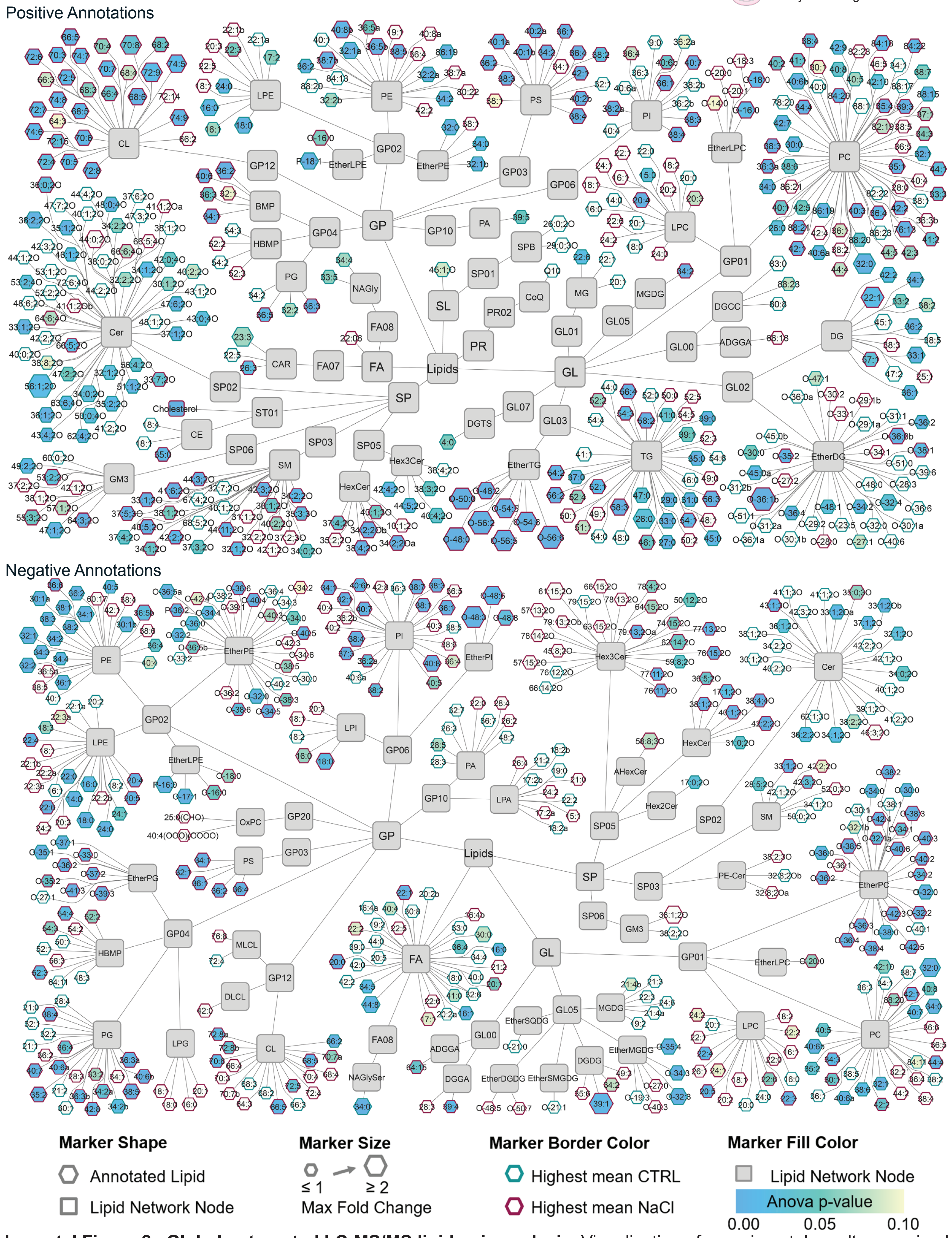


**Supplemental Figure 2. Global untargeted LC-MS/MS lipidomic analysis.** Visualization of experimental results organized in a lipid network by structural similarity. Lipid network nodes are gray rounded rectangles. Lipids detected in positive mode (top) and negative mode (bottom) for one pairwise comparison are represented by terminal hexagons. Terminal hexagon labels specify the annotation, marker size denotes max fold change, border color describes the experimental group with the highest mean, and fill color indicates significance. Annotations observed at two distinct retention times are denoted with “a” for the earlier retention time and “b” for the later retention time.

Supplemental Figure 3

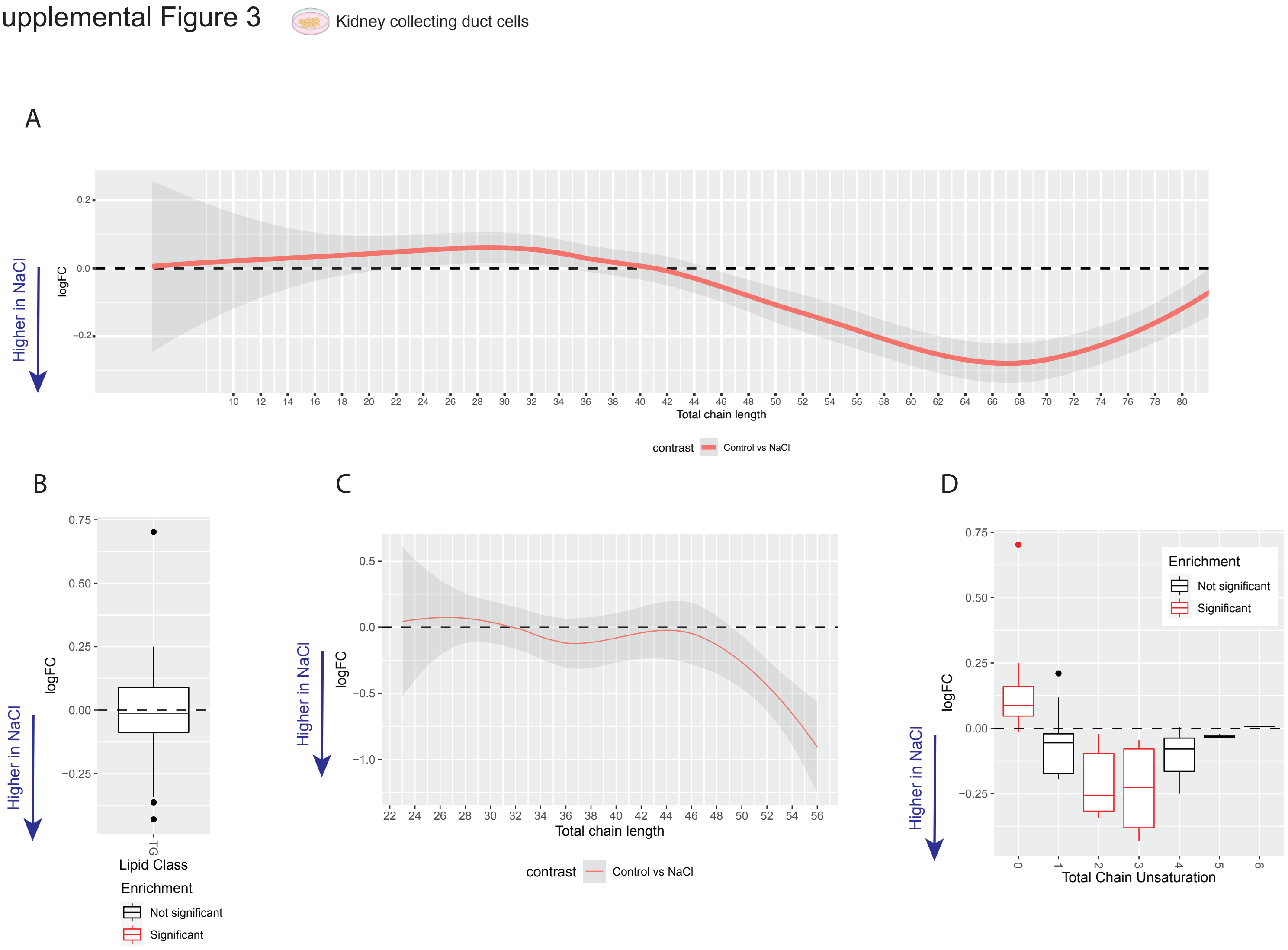


**Supplemental Figure 3. Fatty acid chain length enrichment in kidney collecting duct cells treated with NaCl hypertonic stress vs controls.** (A) Whole cell lipid analysis by total chain length plotted as LogFC enrichment vs total chain length. (B) Analyzed as a lipid class, triglycerides (TG) of all lengths are not significantly enriched in NaCl treated cells. When analyzed by total chain length (C) and unsaturation (D), longer chain unsaturated triglycerides are progressively enriched in NaCl treated cells.

Supplemental Figure 4

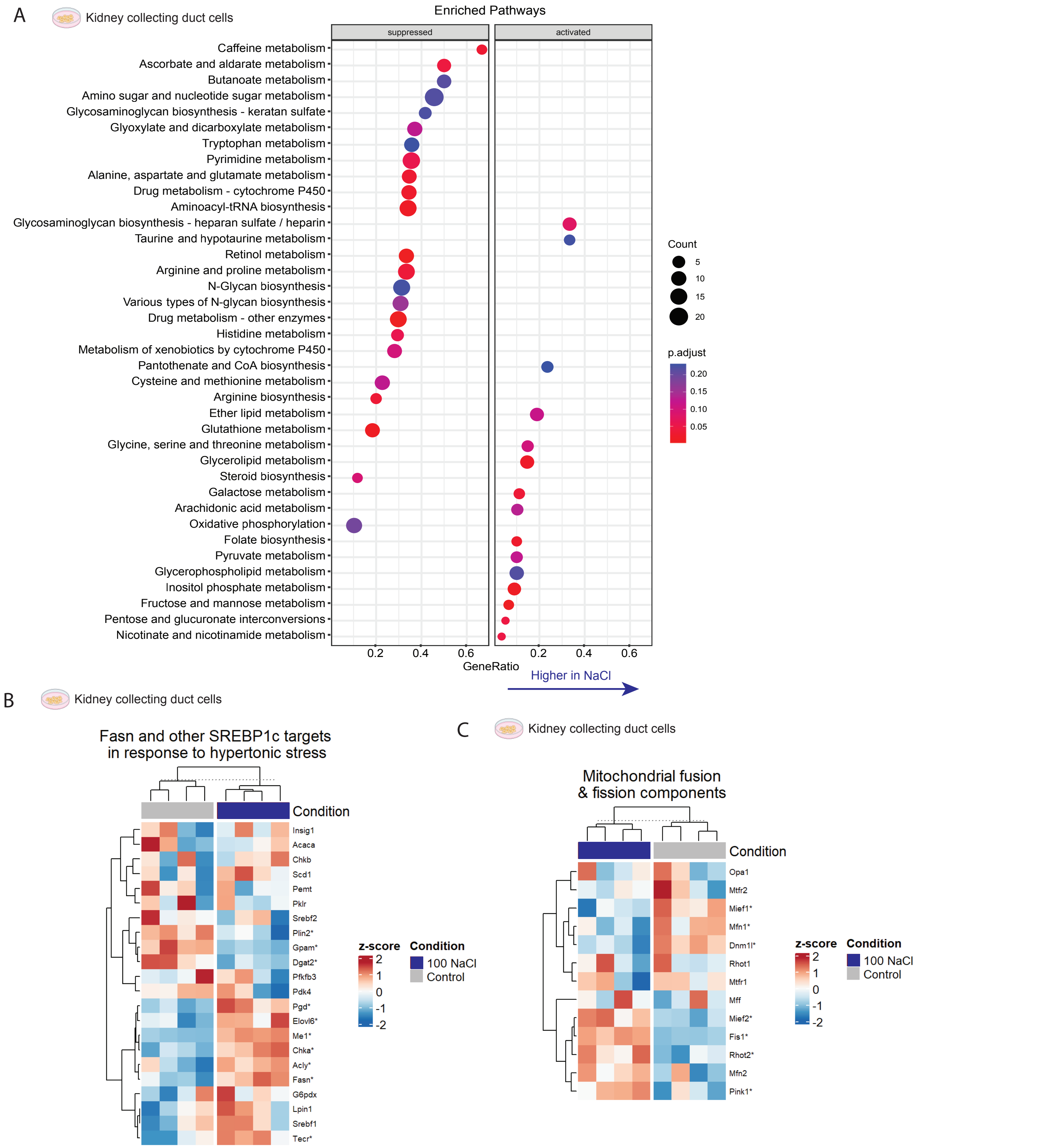


**Supplemental Figure 4. RNA sequencing in kidney collecting duct cells treated with NaCl hypertonic stress vs controls.** (A) Kyoto Encyclopedia of Genes and Genomes (KEGG) pathway analysis from RNA sequencing data from kidney collecting duct cells treated with hypertonic stress vs controls. (B) sterol regulatory element-binding protein 1c (SREBP-1c) target and mitochondrial fusion & fission components enrichment analysis (C) in kidney collecting duct cells treated with hypertonic stress vs controls. * denotes significances with adjusted P value < 0.05

Supplemental Figure 5

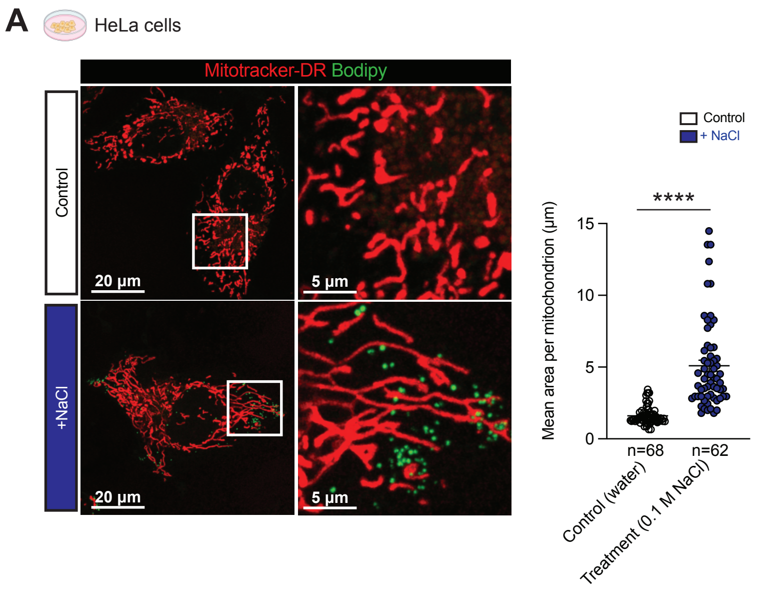


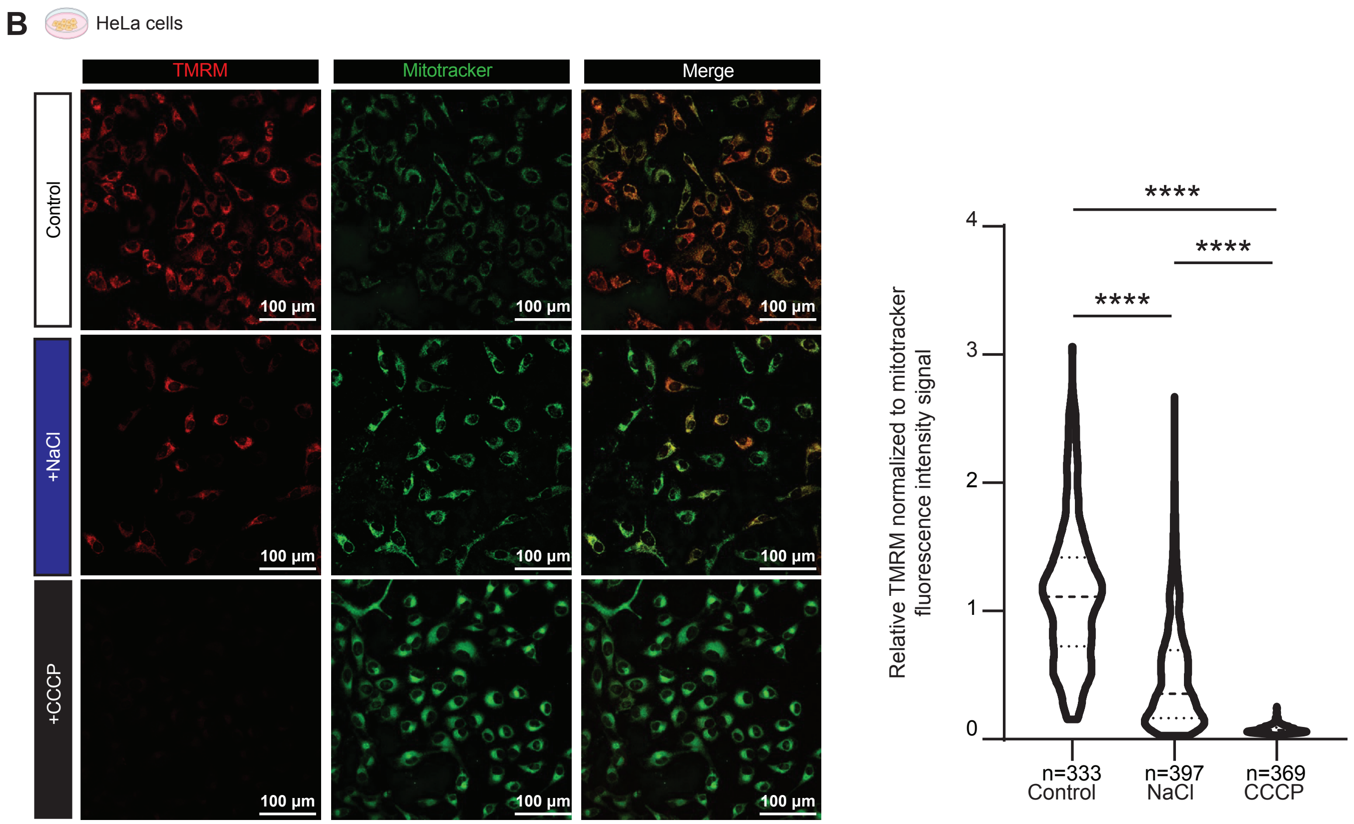


**Supplemental Figure 5 - NaCl challenge alters mitochondrial morphology and function in HeLa cells.** (A) HeLa cells treated with NaCl or control stained with Mitotracker-DR for mitochondrial morphology analysis and co-stained with BODIPY to visualize lipid droplets. (B) Mitochondrial membrane potential analysis in Mitotracker-DR and TMRM stained HeLa cells treated with control, NaCl, or CCCP (carbonyl cyanide m-chlorophenyl hydrazone). Data in A presented as mean +/- SEM, analyzed with a Kruskal-Wallis Test ****<0.0001. Data in B presented as a Violin plot where the median is indicated by a dashed line and the quartiles are indicated by dotted lines, analyzed with a one-way ANOVA ****<0.0001. n as indicated in each panel.

Supplemental Figure 6 - AMPK siNFAT5

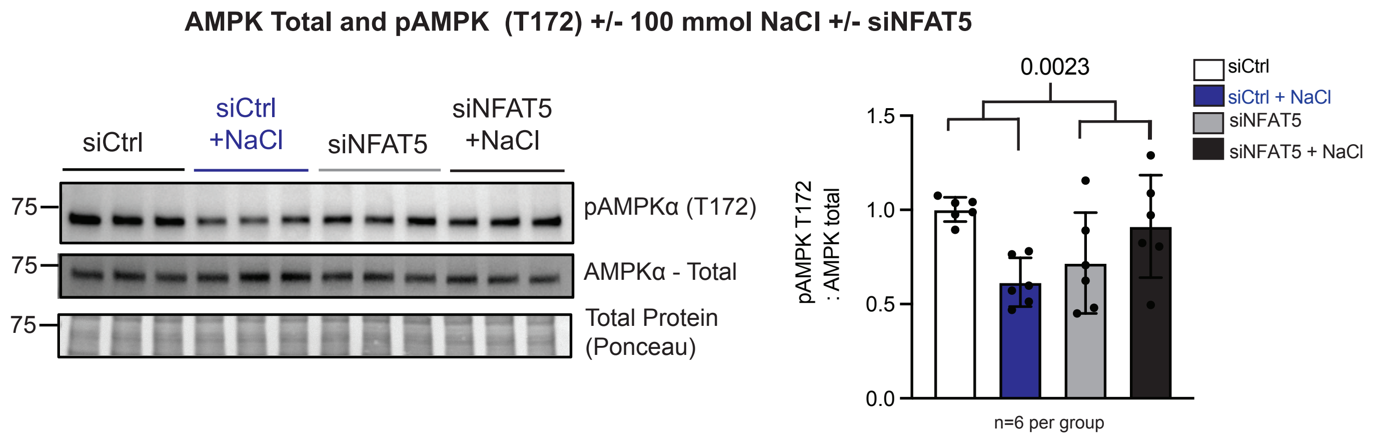


**Supplemental Figure 6. NFAT5 is required for regulation of AMPK during hypertonic stress.** Immunoblots of total AMP activated protein kinase (AMPK) and pAMPKaT172) in kidney collecting duct cells transfected with siNfat5 vs siCtrl treated with NaCl or control.Data presented as mean +/- SD, analyzed with a two-way ANOVA with NaCl and siRNA as the independent variables. p=0.0023 for the interaction, siCtrl vs siCtrl +NaCl p=0.0075, siNFAT5 vs siNFAT5 + NaCl p=0.2152 after Šídák's multiple comparisons test. n as noted in figure.

Supplemental Figure 7-

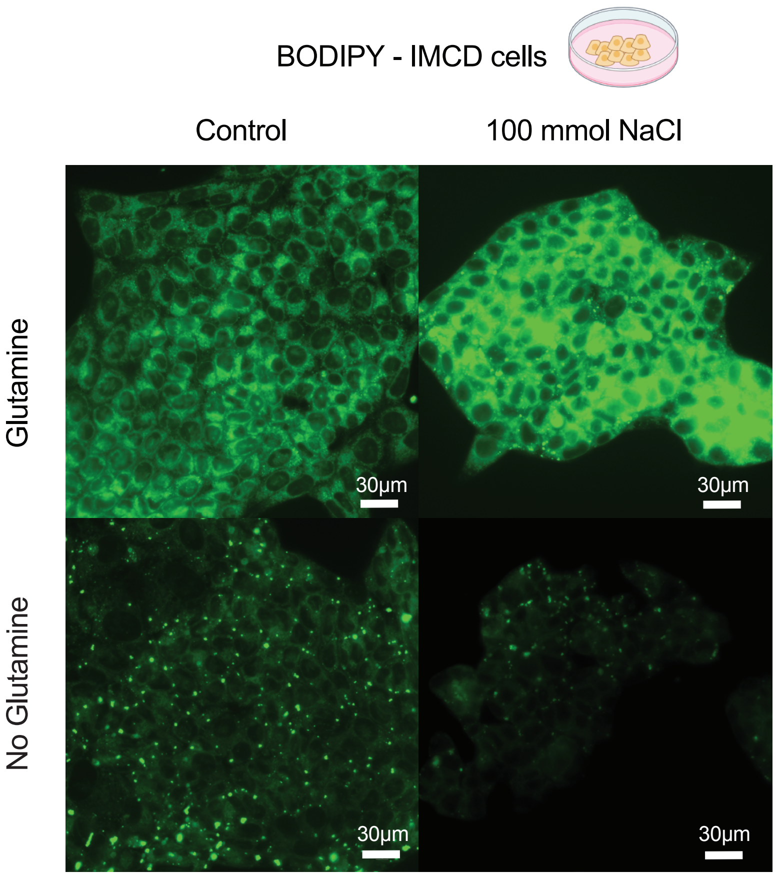


**Supplemental Figure 7 – Effect of glutamine restriction on BODIPY lipid staining in kidney collecting cells** – Intracellular lipids in kidney collecting duct cells treated with NaCl or control in the presence or absence of glutamine in the medium were stained with BODIPY. Representative images shown.

Supplemental Figure 8 - *Drosophila* feeding paradigms

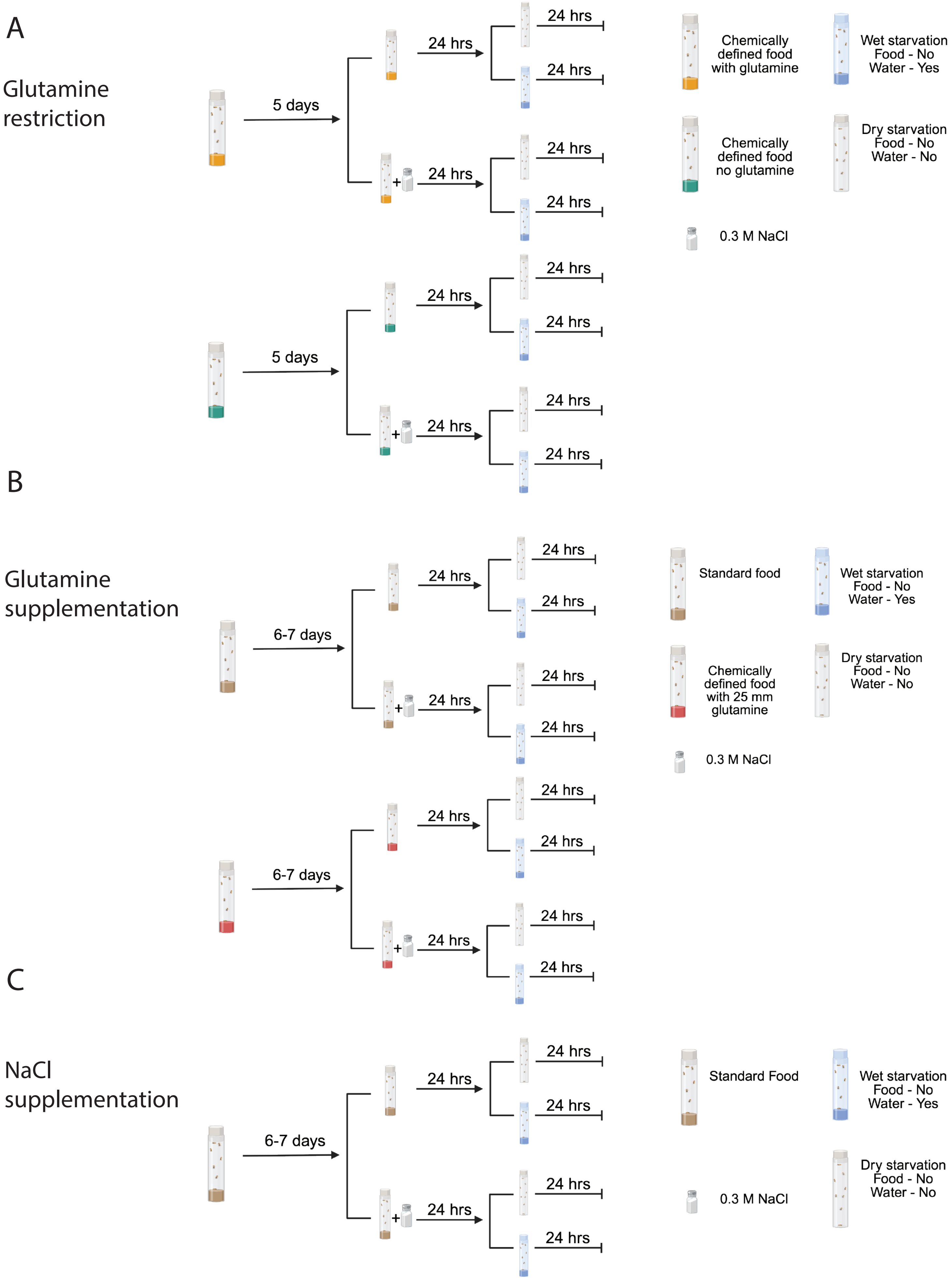

Supplemental Figure 9

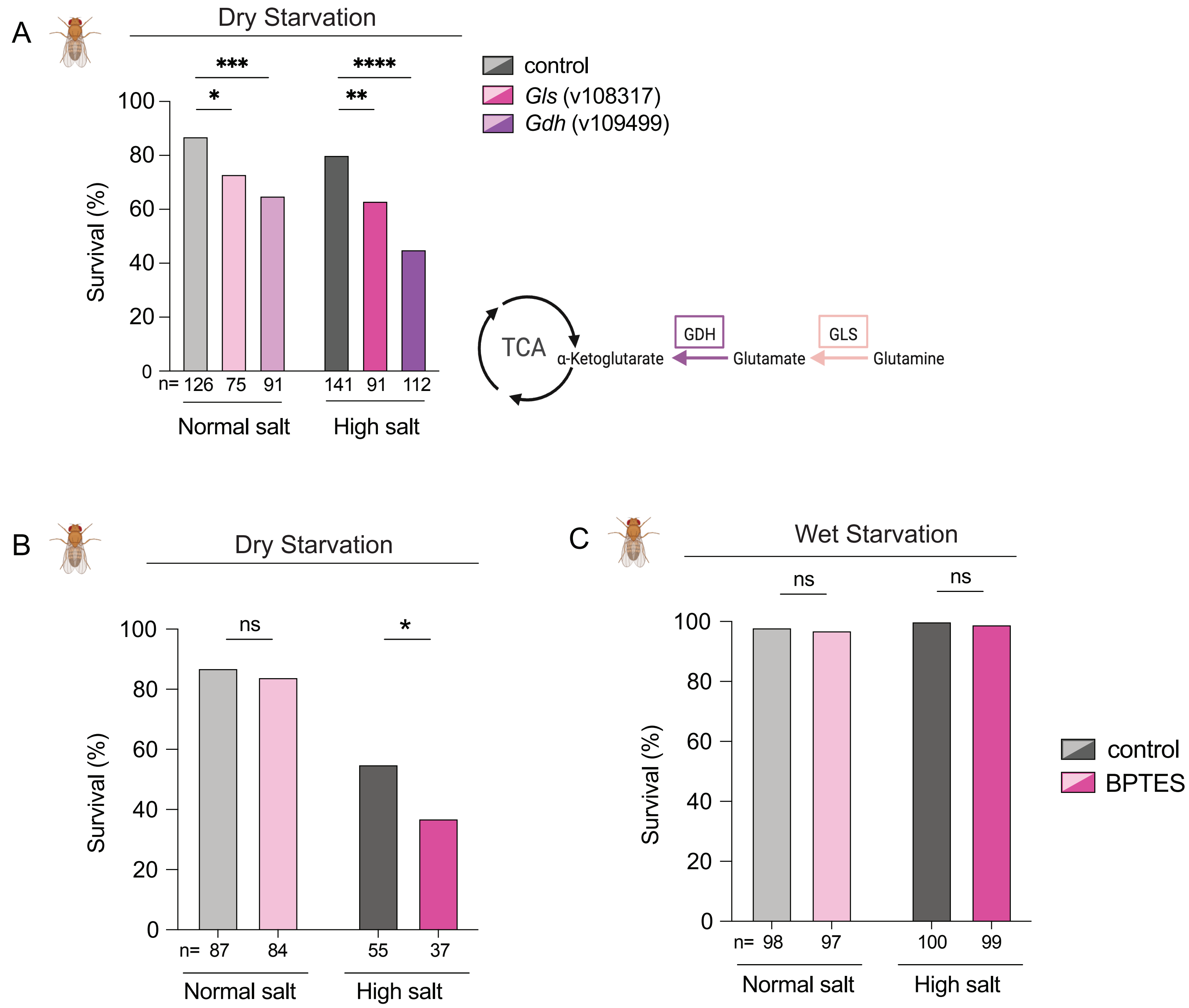


**Supplemental Figure 9 – Additional Gls/Gdh knockdown flies and effects of GLS inhibitor, BPTES** – A) Survival analysis of D. melanogaster with Gls (v108317) and Gdh (v109499) knockdowns (1st set shown in Fig. 4I) when exposed to NaCl hypertonic stress followed by water restriction. B) Survival analysis of D. melanogaster fed DMSO or glutaminase inhibitor BPTES under dry (B) or wet (C) starvation. Data in A,B, and C presented as % survival analyzed with Fischer's exact test. *p<0.05, **p<0.01, ***p<0.001, ****p<0.0001. n as indicated in each panel.

Supplemental Figure 10

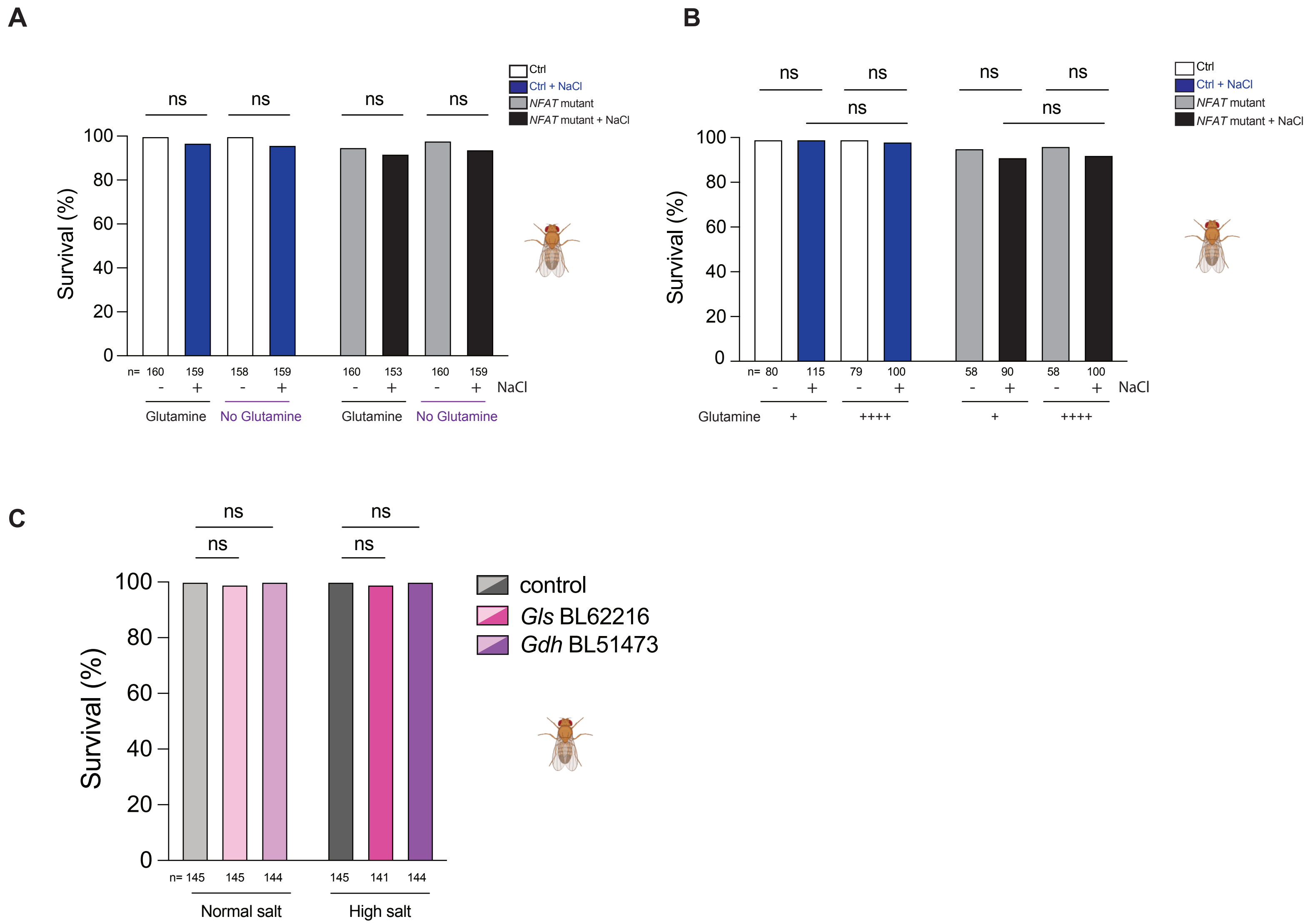


**Supplemental Figure 10 – Wet starvation has no effect on *D. melanogaster* mortality** – Survival analysis in D. melanogaster NFAT mutants vs controls exposed to (A) high salt stress, +/- glutamine, or (B) with supplemented glutamine. (C) Gls (BL62216) and Gdh (BL51473) knockdown flies +/- high salt followed by wet starvation, Data presented as % survival analyzed with Fischer’s exact test. ns = no significant difference. n as indicated in each panel.

| 1ry Antibodies | | | | | |
|---|---|---|---|---|---|
| **Antibody Target** | **Host Species** | **Catalog Number** | **Vendor** | **Dilution and application** | **Time / Temp** |
| ACC total | rabbit | 3662 | Cell Signaling Technologies | WB 1:1000 | 1 hr RT |
| pACC (S79/221) | rabbit | 3661 | Cell Signaling Technologies | WB 1:1000 IF 1:250 | 1 hr RT / 4 C |
| AMPKα (D5A2) | rabbit | 5831 | Cell Signaling Technologies | WB 1:1000 IF 1:250 | 2 hr RT |
| p-AMPKα (Thr172) (40H9) | rabbit | 2535 | Cell Signaling Technologies | WB 1:1000 IF 1:250 | 3 hr RT |
| AQP2 total - FITC conjugated | mouse | sc-515770 | Santa Cruz Biotech | IF 1:100, WB 1:500 | 1 hr RT |
| NFAT5 | rabbit | PA1-023 | Thermo Fisher Scientific | WB 1:1000 | ON 4C |
| pPDH (S293) | rabbit | 31866 | Cell Signaling Technologies | WB 1:1000 | 1 hr RT |
| PDH Total | rabbit | 2784 | Cell Signaling Technologies | WB 1:1000 | 1 hr RT |

| 2ry Antibodies | | | | | |
|---|---|---|---|---|---|
| **Antibody Target** | **Host Species** | **Catalog Number** | **Vendor** | **Dilution and application** | |
| anti-rabbit IgG - HRP conjugated | Goat | 111-035-003 | Jackson Immunoresearch | WB 1:5000 | 1 hr RT |
| anti-rabbit IgG AF 647 conjugated | Donkey | 711-605-152 | Jackson Immunoresearch | IF 1:250 | 1 hr RT |

**Supplemental Table 1 - Antibodies**
